# Interactions of pharmaceutical companies with world countries, cancers and rare diseases from Wikipedia network analysis

**DOI:** 10.1101/614016

**Authors:** Guillaume Rollin, José Lages, Tatiana S. Serebriyskaya, Dima L. Shepelyansky

## Abstract

Using the English Wikipedia network of more than 5 million articles we analyze interactions and interlinks between the 34 largest pharmaceutical companies, 195 world countries, 47 rare renal diseases and 37 types of cancer. The recently developed algorithm using a reduced Google matrix (REGOMAX) allows us to take account both of direct Markov transitions between these articles and also of indirect transitions generated by the pathways between them via the global Wikipedia network. This approach therefore provides a compact description of interactions between these articles that allows us to determine the friendship networks between them, as well as the PageRank sensitivity of countries to pharmaceutical companies and rare renal diseases. We also show that the top pharmaceutical companies in terms of their Wikipedia PageRank are not those with the highest market capitalization.

## Introduction

The improvement of human health and the treatment of various diseases is a vital task for human society [1]. The creation of efficient medications and drugs is now mainly controlled by large biotechnology and pharmaceutical companies. The world’s 34 largest companies are listed in Wikipedia [2]. The analysis of interactions between these companies and their influence on various diseases is an important, but not an easy task. Here, we develop a data mining approach to this task by using the directed network of articles in the English Wikipedia, dated at May 2017, that is generated by citation links between articles. At present, Wikipedia is a repository of a huge amount of human knowledge, exceeding that of the Encyclopedia Britannica in terms of the volume and accuracy of articles devoted to scientific topics [3]. Scientific articles are actively maintained, as it can be seen by the example of articles on biomolecules [4]. Academic research into and analysis of the information present in Wikipedia are growing as new tools and methods are developed, as reviewed in [5]. The quality of Wikipedia articles is improving with time as shown by the analysis reported in [6].

At present a variety of methods exist for the analysis and characterization of complex networks (see e.g. [7]). The most popular analytical tool is the PageRank algorithm invented by Brin and Page in 1998 for ranking the sites on the World Wide Web (WWW) [8]. A detailed description of the algorithm and the mathematical properties of the related Google matrix are given in [9] while a variety of applications of the Google matrix to real directed networks are described in [10]. The application of Google matrix methods to Wikipedia networks in 24 different languages has led to a reliable ranking of historical figures over 15 centuries of human history [11], as well as to the ranking of the world’s universities [12].

Recently, the reduced Google matrix (REGOMAX) algorithm has been proposed for the analysis of the effective interactions and links between a selected subset of nodes of interest embedded in a much larger global network [13]. The REGOMAX algorithm originates from the scattering theory of nuclear and mesoscopic physics and the field of quantum chaos. Its efficiency has been demonstrated for Wikipedia networks retrieving known effective interactions between politicians [14], geopolitical relations and links between world countries [15], universities [16] and banks [17].

The REGOMAX approach has also been used for networks of protein-protein interactions [18] and for Wikipedia networks in order to determine the influence in Wikipedia of infectious diseases [19], drugs and cancers [20]. Here, we extend this approach to analyze the influence and mutual interactions in Wikipedia of the world’s 34 largest biotechnology and pharmaceutical companies [2]. As the PageRank and the REGOMAX algorithms provide rankings of items according to how these items are connected, an item will, in the following, be designated as influential in Wikipedia if its ranking, mirroring the cascade of directed links converging toward it, is high. We use the Wikipedia network with 5 416 537 articles and establish relations between 195 countries and these companies on the basis of their PageRank sensitivity (see section Methods) of the corresponding articles. We also construct interaction networks between these companies and 47 rare renal diseases and 37 types of cancer. The number of people suffering from cancer in 2012 was about 32.6 million worldwide, with this number increasing by 6-9 million people each year [21]. By 2019, the number of people worldwide suffering from rare diseases had reached 350 million [22]. Despite the fact that cancer and 80% of rare diseases are genetic diseases, the challenge involved in developing drugs for these two groups of diseases differ. One of the fundamental challenge in the field of drug development for rare diseases is the huge number and diversity of such diseases. There are more than 7 000 known rare diseases [22] whereas the number of known cancer types is quite limited. The large number and diversity of rare diseases cause problems at the level of drug development due to the small number of patients accessible for individual disease, the logistics involved in reaching widely dispersed patient populations, the lack of validated biomarkers and surrogate end-points, and the limited clinical expertise and centers of excellence available [23]. The representation of information about cancer and rare diseases in Wikipedia clearly illustrates the difference between cancers and rare diseases. Most of the cancer types listed on the National Cancer Institute site [24] are represented in Wikipedia. At the same time, we observe that only 8.5% and 15% of the phenotypes described in Orphanet [25] and OMIM [26], respectively, are addressed by topics in Wikipedia. For our calculations, we use only the 47 rare renal diseases listed in [27] and represented in Wikipedia.

We hope that, through the prism of Wikipedia, our results will make it possible to gain a better understanding of the influence of pharmaceutical companies on these diseases and to obtain specialization profiles for these pharmaceutical companies.

The paper is constructed as follows: the mathematical methods are described in the Section Methods and data are overviewed in the Section Datasets. The two following Sections are devoted to the presentation of Results and of the Discussion.

## Methods

### Google matrix construction

A detailed description of the construction of the Google matrix and its properties can be found in [9, 10]. Here, we therefore provide only a brief description in order to assist readers.

Let us consider a network of *N* nodes described by the adjacency matrix *A* the elements of which are either *A*_*ij*_ = 1 if node *j* points to node *i*, or *A*_*ij*_ = 0 otherwise. The elements of the Google matrix *G* have the standard form *G*_*ij*_ = *αS*_*ij*_ + (1 *− α*)*/N* [8–10], where the matrix *S* describes Markov transitions from node to node. For a random surfer, the matrix element *S*_*ij*_ gives the probability of transition from node *j* to node *i* according to the directed network structure. The elements of matrix *S* are either *S*_*ij*_ = *A*_*ij*_/*k*_*out*_(*j*) if the out-degree of node *j*, 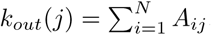, is non zero, or *S*_*ij*_ = 1/*N* if *j* has no outgoing links (dangling node), i.e., *k*_*out*_(*j*) = 0. Hence, the random surfer can jump to any node once it reaches a dangling node. Here *α* is the damping factor (0 *< α <* 1) and we use its standard value *α* = 0.85 [9] in the following. It should be noted that for the range 0.5 ≤ *α* ≤ 0.95, the results are not sensitive to *α* [9, 10]. At each jump, the random surfer either follows the network structure with a transition probability proportional to *α*, or randomly chooses any node with a transition probability proportional to 1 − *α*. The damping factor *α* prevents the random surfer from being trapped in dangling subnetworks [9].

Starting its journey from node *j*_0_, after *n* iterations the random surfer will arrive at node *i* with the probability 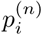, which is given by the *i*th component of the probability vector *p*^(*n*)^ = *G*^*n*^*p*^(0)^ where 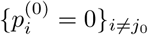 and 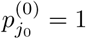. According to the Perron-Frobenius theorem [9], for any initial vector *p*^(0)^, the iteration process converges to a unique vector *P*, which, within the framework of the Google matrix is called the PageRank vector, and is given by the following limit *P* = lim_*n→∞*_ *G*^n^*v* for any vector *v* with positive elements and 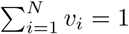. Consequently, the PageRank vector *P* is the steady state vector, *P* = *GP*, of the Markovian process describing the endless journey of the random surfer. Furthermore, the PageRank vector is the right eigenvector of the Google matrix *G* associated with the eigenvalue 1. The *i*th component of the PageRank vector *P* gives the probability of finding the random surfer at node *i* after its endless journey, or, in practical terms, after a sufficiently high number of iterations in order to ensure the convergence toward the PageRank vector *P*. We order all the nodes initially labeled by *i* = 1, …, *N* by decreasing PageRank probability {*P*_*j*_}_*j=1,…,N*_ introducing PageRank indexes *K* = 1, …, *N*. A node *j* is ranked higher than a node *i*, i.e. *K*(*j*) *< K*(*i*), if *P_j_ > P*_*i*_. The index *K* = 1 is then assigned to the node with the highest PageRank probability, the index *K* = 2 is assigned to the node with the second highest PageRank probability, and so on. The index *K* = *N* is assigned to the node with the lowest PageRank probability. The numerical computation of PageRank probabilities is performed efficiently using the PageRank iteration algorithm described in [8, 9].

It is also useful to consider the network obtained by reversing the direction of all directed links of the original network. With this reversed network, we associate the Google matrix *G^∗^*, which is built from the adjacency matrix *A^∗^* = *A*^t^ which is the transpose of the original adjacency matrix *A*. The elements of the stochastic matrix *S^∗^* are either 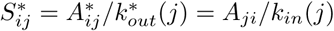, if 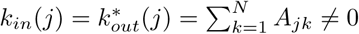, or 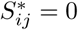, otherwise. The elements of the Google matrix *G^∗^* are 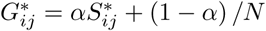. The PageRank vector associated with *G^∗^* is called the CheiRank vector *P ^∗^*. It characterizes the steady state, *P ^∗^* = *G^∗^P ^∗^*, of the Markovian process described by the matrix *G*^*∗*^ [28, 29] (see also [10]). Its components, 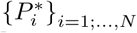, are also probabilities that can again be ordered by decreasing values. We introduce the CheiRank indexes *K*^*∗*^ = 1, …, *N* in such a way that we assign *K*^*∗*^ = 1 (*K*^*∗*^ = *N*) to the node with the highest (lowest) CheiRank probability. On average, high PageRank (CheiRank) probabilities correspond to nodes with many ingoing (outgoing) links [9, 10].

The PageRank measure of centrality characterizes the influence of a node inside the network since it quantifies how other nodes point toward it. By contrast, the CheiRank measure of centrality characterizes the diffusivity of a node since it quantifies how it points toward the other nodes of the network. To summarize, according to the PageRank algorithm, a node is all the more influential if it is pointed to by influential nodes while, according to the CheiRank algorithm, a node is all the more diffusive if it points to diffusive nodes.

### Reduced Google matrix

The REGOMAX algorithm was proposed in [13] and is described in detail in [14–16, 18]. Here, we present the main elements of this method using the notation employed in [14].

Let us consider a subset *S*_*r*_ of *N*_*r*_ nodes of interest from a huge network of *N ≫ N_r_* nodes. The REGOMAX algorithm efficiently computes a *reduced Google matrix G*_R_ of size *N_r_ × N*_*r*_ that captures the full contributions of direct and indirect pathways between any pair of nodes among the *N*_*r*_ nodes of interest. For a random surfer navigating the entire network during an infinite journey, the reduced Google matrix entry *G*_R*ij*_ gives the probability of transition from node *j* to node *i* (*i, j ∈S*_*r*_), while taking into account the probability of a possible direct transition from node *j* to node *i*, and also taking account of the indirect transition probabilities for all the indirect pathways from node *j* to node *i* via at least one node *k* not belonging to the nodes of interest subset *S*_*r*_. Mathematically, the reduced Google matrix is unequivocally determined once we assume that the relative PageRank probabilities of the *N*_*r*_ nodes obtained from the *N_r_ × N*_*r*_ reduced Google matrix *G*_R_ are the same as those obtained from the global *N × N* Google matrix *G*. The reduced Google matrix *G*_R_ can be decomposed into three matrix components that clearly distinguish between direct and indirect interactions, *G*_R_ = *G*_*rr*_ + *G*_pr_ + *G*_qr_ [14]. Here, the *G*_*rr*_ matrix component is a *N_r_ × N*_*r*_ submatrix of the global *N × N* Google matrix *G* corresponding to the direct transition probabilities between the *N*_*r*_ nodes of interest. The *G*_pr_ matrix component is quite similar to the matrix in which each column is equal to the PageRank vector *P*_*r*_, which is defined as *G*_R_*P*_*r*_ = *P*_*r*_. As the PageRank vector components 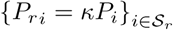 obtained from *G*_R_ are straightforwardly deduced from the PageRank vector components 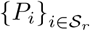, with the coefficient *κ* being a constant for all *i*, the *G*_pr_ matrix component does not provide much more information than the global PageRank vector *P* about direct and indirect links between the *N*_*r*_ selected nodes. Thus the most interesting and nontrivial role is played by the *G*_qr_ matrix component. This matrix component takes account of all indirect links between the *N*_*r*_ selected nodes emerging from myriads of pathways passing through multiple nodes of the global network but not belonging to the nodes of interest subset *S*_*r*_ (see [13, 14]). We decompose the *G*_qr_ = *G*_qrd_ + *G*_qrnd_ matrix component into its diagonal part (*G*_qrd_) and its non-diagonal part (*G*_qrnd_). The *G*_qrd_ diagonal matrix component contains self-interactions of nodes which are not of interest for the present study. The precise formulas used to compute the three matrix components of the reduced Google matrix *G*_R_ are given in [13, 14].

A useful additional characteristic provided by the *G*_R_ matrix is the sensitivity of the PageRank probability to the variation of a specific link between a pair of nodes chosen among the *N*_*r*_ nodes of interest. The useful results obtained with this method have been demonstrated in [15–17, 19, 20]. The PageRank sensitivity *D*(*j → k, i*) of a node *i* to the matrix element *G*_R*jk*_ is

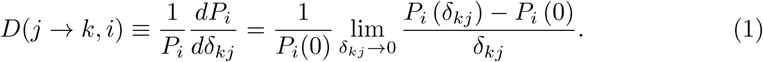

Here, *P* (*δ*_*kj*_) is the PageRank vector obtained from the reduced Google matrix *G*_R_, where all the elements of *j*th column are modified as 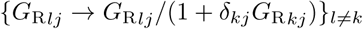, with the exception of the *k*th element which is transformed as 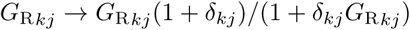. This transformation ensures that the sum of these transformed elements stays equal to 1, 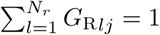. This is a mandatory property since the elements of the *j*th column of *G*_R_ represent the probabilities of transitions from node *j* to the other nodes. Roughly speaking, forgetting the denominator 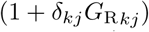, which ensures the normalization of the *j*th column, the PageRank sensitivity *D*(*j → k, i*) measures the relative variation of the PageRank of the *i*th node when the *G*_R_ link from node *j* to node *k* changes by an infinitesimal amount.

### Datasets

We consider the English Wikipedia edition as at May 2017 with *N* = 5 416 537 articles (nodes) and *N*_*l*_ = 122 232 932 hyperlinks between articles. This network has also been considered in [16, 17, 19, 20]. For the REGOMAX analysis, we select *N*_*c*_ = 195 world countries (see the list and PageRank order in [19, 20]), the *N*_*ph*_ = 34 largest pharmaceutical companies (see Table 1), *N*_*rd*_ = 47 rare renal diseases (see Table 2), and *N*_*cr*_ = 37 types of cancer listed in [20]. Thus we consider a total of *N*_*r*_ = *N*_*c*_ + *N*_*ph*_ + *N*_*rd*_ + *N*_*cr*_ = 313 Wikipedia articles as nodes of interest. All data sets are available at [30]. The present study complies with the Wikimedia terms of use.

**Table 1.** List of the 34 largest pharmaceutical companies ranked by the relative PageRank index *K*_*r*_ of their corresponding articles in Wikipedia. The *K*_*LMC*_ index gives the ranking by the largest market capitalization since 2000 [2] and the *K*_*MC*_ index gives the ranking by market capitalization in 2017 [2]. The relative CheiRank index 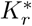 is also given.

| $K_r$ | $K_r^*$ | $K_{LMC}$ | $K_{MC}$ | Company | $K_r$ | $K_r^*$ | $K_{LMC}$ | $K_{MC}$ | Company |
| --- | --- | --- | --- | --- | --- | --- | --- | --- | --- |
| 1 | 3 | 2 | 3 | Pfizer | 18 | 16 | 18 | 18 | Biogen |
| 2 | 1 | 8 | 15 | GSK<sup>a</sup> | 19 | 25 | 28 | 28 | Mylan |
| 3 | 8 | 14 | 10 | Bayer | 20 | 13 | 7 | 13 | Gilead |
| 4 | 10 | 1 | 1 | J&J<sup>b</sup> | 21 | 17 | 21 | 21 | Shire |
| 5 | 2 | 3 | 4 | Novartis | 22 | 22 | 23 | 23 | Takeda |
| 6 | 4 | 6 | 6 | Merck | 23 | 28 | 29 | 27 | Astellas |
| 7 | 6 | 16 | 14 | Lilly | 24 | 18 | 32 | 29 | Daiichi Sankyo |
| 8 | 11 | 4 | 2 | Roche | 25 | 15 | 5 | 5 | AbbVie |
| 9 | 5 | 19 | 16 | AstraZeneca | 26 | 32 | 22 | 22 | Regeneron |
| 10 | 12 | 10 | 9 | Sanofi | 27 | 24 | 33 | 31 | Eisai |
| 11 | 9 | 15 | 11 | BMS<sup>c</sup> | 28 | 31 | 20 | 19 | Stryker |
| 12 | 7 | 13 | 12 | Abbott | 29 | 19 | 34 | 33 | BioMarin |
| 13 | 29 | 24 | 24 | Illumina | 30 | 26 | 27 | 26 | Zoetis |
| 14 | 14 | 11 | 8 | Amgen | 31 | 23 | 25 | 25 | Vertex |
| 15 | 21 | 9 | 7 | Novo Nordisk | 32 | 30 | 26 | 30 | Alexion |
| 16 | 33 | 12 | 20 | Allergan | 33 | 34 | 31 | 34 | Perrigo |
| 17 | 20 | 17 | 17 | Celgene | 34 | 27 | 30 | 32 | Incyte |

<sup>a</sup>GSK: GlaxoSmithKline, <sup>b</sup>J&J: Johnson & Johnson, <sup>c</sup>BMS: Bristol-Myers Squibb.

**Table 2.** List of 47 rare renal diseases ranked by the relative PageRank index *K*_*r*_ of their corresponding articles in Wikipedia. The relative CheiRank index, 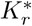, is also given. The list of rare renal diseases is split in 5 categories with the following color code: 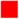 Congenital abnormalities of the kidney and urinary tract, 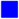 Glomerular diseases, 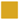 Renal tubular diseases and metabolic diseases, 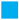 Nephrolithiasis and 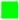 Ciliopathies and Nephronophthisis.

| $K_r$ | $K_r^*$ | | Rare renal disease | Short name |
| --- | --- | --- | --- | --- |
| 1 | 6 | ■ | Alport syndrome | Alport |
| 2 | 1 | ■ | Bardet–Biedl syndrome | Bardet–Biedl |
| 3 | 7 | ■ | Fabry disease | Fabry |
| 4 | 2 | ■ | Kallmann syndrome | KS |
| 5 | 4 | ■ | Renal tubular acidosis | RTA |
| 6 | 5 | ■ | Cystinuria | Cystinuria |
| 7 | 14 | ■ | Renal agenesis | Renal agenesis |
| 8 | 16 | ■ | Nephronophthisis | Nephronophthisis |
| 9 | 33 | ■ | X-linked hypophosphatemia | XLH |
| 10 | 8 | ■ | Bartter syndrome | Bartter |
| 11 | 41 | ■ | Xanthinuria | Xanthinuria |
| 12 | 3 | ■ | Oculocerebrorenal syndrome | Oculocerebrorenal |
| 13 | 17 | ■ | Nail–patella syndrome | Nail–patella |
| 14 | 45 | ■ | Familial renal amyloidosis | Amyloidosis |
| 15 | 9 | ■ | Episodic ataxia | EA |
| 16 | 24 | ■ | Gitelman syndrome | Gitelman |
| 17 | 21 | ■ | Medullary cystic kidney disease | MCKD |
| 18 | 32 | ■ | Familial hypocalciuric hypercalcemia | FHH |
| 19 | 42 | ■ | Renal glycosuria | Glycosuria |
| 20 | 40 | ■ | Fanconi–Bickel syndrome | Fanconi–Bickel |
| 21 | 23 | ■ | Liddle’s syndrome | Liddle |
| 22 | 31 | ■ | Hypomagnesemia with secondary hypocalcemia | HSH |
| 23 | 13 | ■ | Fraser syndrome | Fraser |
| 24 | 36 | ■ | WAGR syndrome | WAGR |
| 25 | 19 | ■ | Branchio–oto–renal syndrome | BOR |
| 26 | 26 | ■ | Townes–Brocks syndrome | Townes–Brocks |
| 27 | 44 | ■ | Autosomal dominant hypophosphatemic rickets | ADHR |
| 28 | 18 | ■ | Orofaciodigital syndrome 1 | Orofaciodigital |
| 29 | 28 | ■ | Denys–Drash syndrome | DDS |
| 30 | 10 | ■ | Perlman syndrome | Perlman |
| 31 | 11 | ■ | Simpson–Golabi–Behmel syndrome | SGBS |
| 32 | 12 | ■ | Caroli disease | Caroli |
| 33 | 25 | ■ | Congenital nephrotic syndrome | GNS |
| 34 | 39 | ■ | Fechtner syndrome | Fechtner |
| 35 | 27 | ■ | Abderhalden–Kaufmann–Lignac syndrome | AKL |
| 36 | 22 | ■ | Lysinuric protein intolerance | LPI |
| 37 | 47 | ■ | Majewski’s polydactyly syndrome | Majewski |
| 38 | 20 | ■ | Urofacial syndrome | Urofacial |
| 39 | 15 | ■ | Barakat syndrome | Barakat |
| 40 | 35 | ■ | Dicarboxylic aminoaciduria | DCBXA |
| 41 | 38 | ■ | MODY 5 | MODY 5 |
| 42 | 30 | ■ | Papillorenal syndrome | Papillorenal |
| 43 | 43 | ■ | OCRL | OCRL |
| 44 | 37 | ■ | COQ9 | COQ9 |
| 45 | 29 | ■ | Juvenile nephronophthisis | J. nephronophthisis |
| 46 | 46 | ■ | Renal–hepatic–pancreatic dysplasia | RHPD |
| 47 | 34 | ■ | Conorenal syndrome | Conorenal |

In Table 2, the 47 rare renal diseases can be grouped into five categories with which we have associated a color to simplify identification: congenital abnormalities of the kidney and urinary tract (red), glomerular diseases (blue), renal tubular diseases and metabolic diseases (gold), nephrolithiasis (cyan), and ciliopathies and nephronophthisis (green). This reflects the categorization used in [27]. Of the 166 genetic disorders of renal growth and of renal function listed in [27], only the 47 rare renal diseases cited above had a corresponding article in Wikipedia dated at May 2017.

## Results

### PageRank vs CheiRank distributions

In Fig 1 we show the coarse-grained distribution of all *N* Wikipedia articles on the PageRank-CheiRank plane (*K, K*^*∗*^) in logarithmic scale. On this plane, we plot the positions of 195 countries (white circles), 37 cancer types (green circles), 47 rare renal diseases (gold circles), 34 pharmaceutical companies (purple circles) and we also add the 230 infectious diseases (red circles) studied in [19]. As usual, countries take the top PageRank positions (see also [10]), followed by the groups of infectious diseases and cancers. The group of rare diseases is clustered in the high *K* and high *K*^*∗*^ region, being generally far lower-ranked. The highest-ranked company in the group of pharmaceutical companies is a little behind the leader of the group of cancers. In the global PageRank list the top 3 in each group are *K* = 1 US, *K* = 2 France, *K* = 3 Germany for countries; *K* = 639 tuberculosis, *K* = 810 HIV/AIDS, *K* = 1116 malaria for infectious diseases [19]; *K* = 3478 lung cancer, *K* = 3788 breast cancer, *K* = 3871 leukemia for cancers [20]; *K* = 9345 Pfizer, *K* = 11290 GlaxoSmithKline, *K* = 13737 Bayer for pharmaceutical companies, *K* = 156963 Alport syndrome, *K* = 161731 Bardet–Biedl syndrome, *K* = 174780 Fabry disease for rare renal diseases.

**Fig 1.**
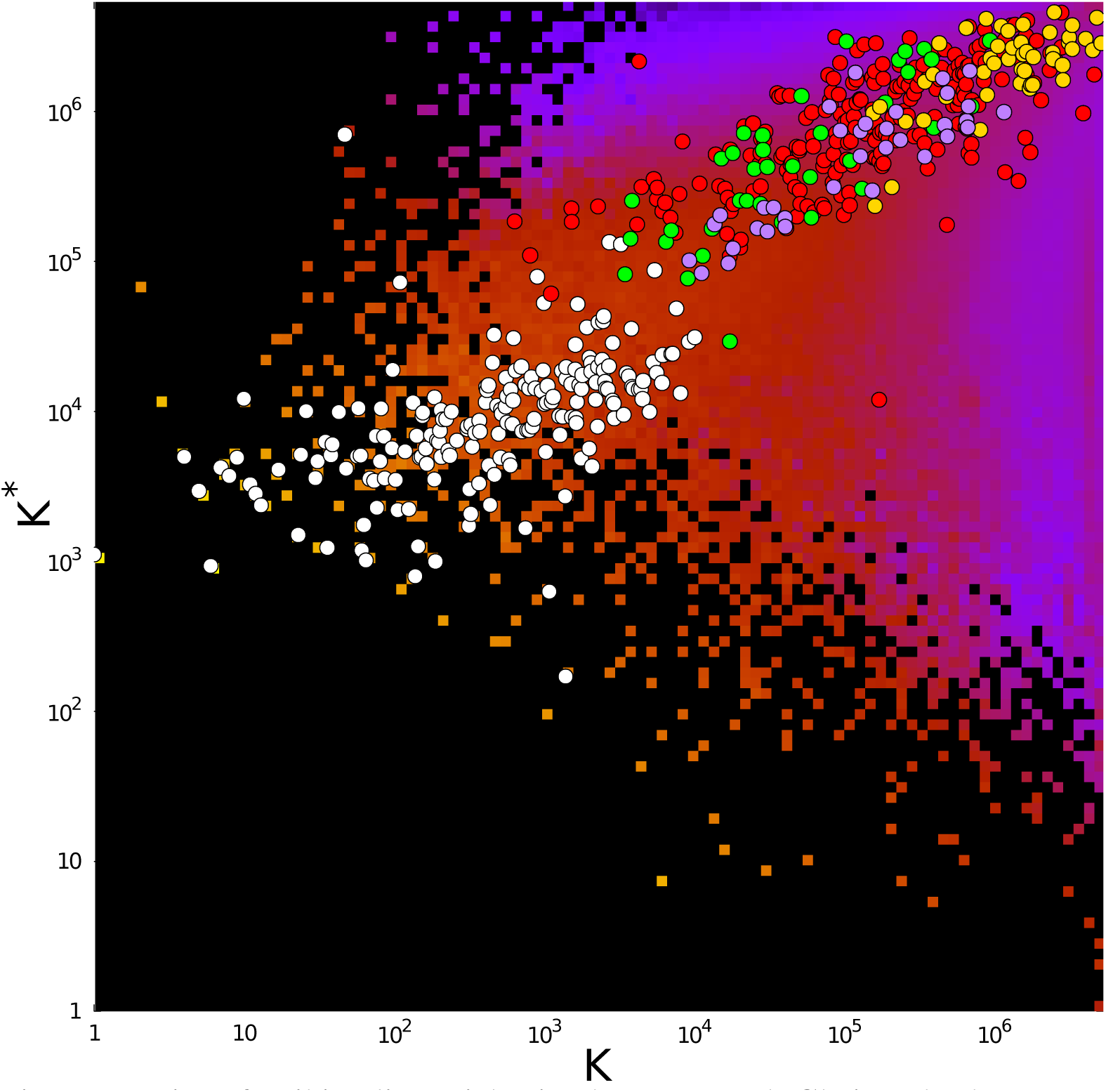
Density of Wikipedia articles in the PageRank-CheiRank plane (*K*, *K*^∗^). Data are averaged over a 100 *×* 100 grid for (log_10_ *K*, log_10_ *K*^*∗*^) spanning the domain [0, log_10_ *N*] × [0, log_10_ *N*]. Density of articles ranges from very low (purple tiles) to very high (bright yellow tiles). The absence of any article is represented by black tiles. The superimposed white circles give the positions of articles devoted to the 195 countries, the red circles represent the positions of articles devoted to the 230 kinds of infectious diseases studied in [19], the green circles represent the positions of 37 articles on the cancers studied in [20], the gold circles mark the positions of 47 articles on rare renal diseases and the purple circles give the positions of 34 articles on pharmaceutical companies.

In Table 1, we give the relative PageRank *K*_*r*_ list of 34 pharmaceutical companies. In this table, we also indicate the rank *K*_*LMC*_ of these companies in terms of largest market capitalization [2] and the rank *K*_*MC*_ based on their market capitalization ranking in 2017 [2]. The CheiRank rank 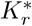 is also given. To permit a quick comparison, Table 3 presents the Spearman rank correlation coefficients *ρ* between these rankings. As expected, the *K*_*LMC*_ and *K*_*MC*_ rankings are similar being highly correlated. We observe that the PageRank *K*_*r*_ and CheiRank 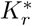 are also highly correlated. Consequently, on average, the influence and the diffusivity of an article on a pharmaceutical company in Wikipedia are correlated. We also observe a fairly high correlation *∼* 0.8 between PageRank *K*_*r*_ and market capitalization rankings *K*_*MC*_ or *K*_*LMC*_. The influence of pharmaceutical company articles within Wikipedia is consequently related to the real economic performances of the pharmaceutical companies. The correlation between CheiRank 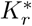 and the *K*_*MC*_ or *K*_*LMC*_ rankings is a little weaker at about *∼* 0.7.

**Table 3.**
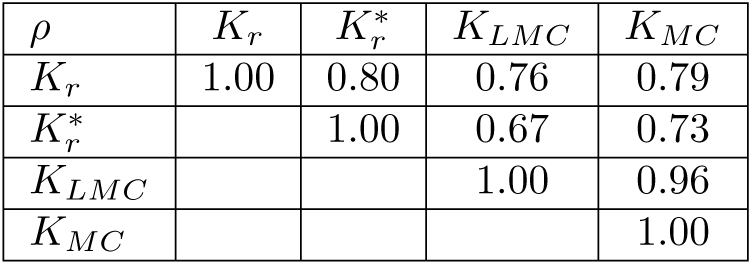
Spearman rank correlation coefficients *ρ* between the *K*_*r*_, 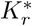, *K*_*LMC*_, and *K*_*MC*_ rankings for pharmaceutical companies presented in Table 1.

The distribution of the companies on the rank plane (*K*_*r*_, *K*_*LMC*_) is shown in Fig 2. If the influence in Wikipedia of the pharmaceutical companies were strictly proportional to their wealth, the companies would lie on the diagonal, *K*_*r*_ = *K*_*LMC*_. Consequently, in this figure, companies above (below) the diagonal have an excess (shortfall) of influence in Wikipedia relative to their market capitalization. The richest company, Johnson & Johnson (J&J), is only at the 4th position in PageRank (*K*_*r*_ = 4). This shows that its influence in Wikipedia is well behind its first place in terms of market capitalization. We can also note that AbbVie, a spin-off of Abbott laboratories, which has enjoyed the 5th largest market capitalization since 2000 is among the less influential of the considered pharmaceutical companies with *K*_*r*_ = 25. In contrast, Pfizer is the most influential company on Wikipedia with *K*_*r*_ = 1 and *K*_*LMC*_ = 2. A significant influence in Wikipedia is found for pharmaceutical companies with a leading *K*_*r*_ position and a somewhat worse *K*_*LMC*_. Such companies include GlaxoSmithKline (GSK) with *K*_*r*_ = 2 and *K*_*LMC*_ = 8 and Bayer with *K*_*r*_ = 3 and *K*_*LMC*_ = 14.

**Fig 2.**
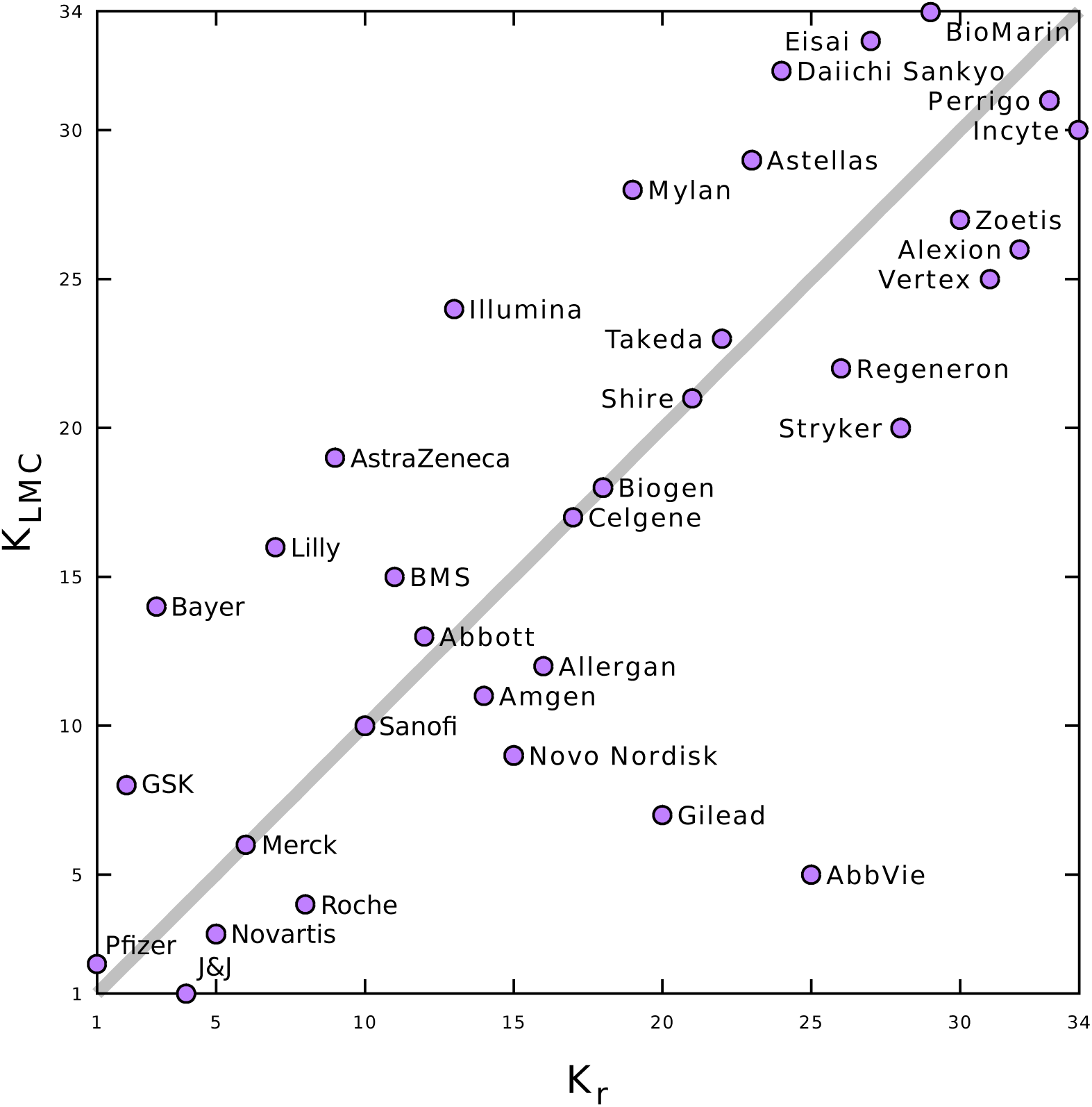
Distribution of pharmaceutical companies ranked by largest market capitalization index, *K*_*LMC*_, and by the relative PageRank index, *K*_*r*_, of their article in Wikipedia. See rankings in Table 1.

The overlap *η*(*j*) between the two ranking lists *K*_*r*_ and *K*_*LMC*_ (see Table 1), as well as between the two ranking lists *K*_*r*_ and *K*_*MC*_ (see Table 1), is shown in Fig 3. We see that for the first three positions, the overlap is only *η* = 1*/*3. Assuming that the major search engines still probe the structural linking properties of the WWW, this shows that the richest pharmaceutical companies can significantly improve their public visibility in Wikipedia. At present, the Wikipedia web site is the 5th most frequently visited site in the world [31] and the improvement in the content of Wikipedia articles can bring significantly better global visibility for companies; furthermore article editing is free of charge. Nevertheless 7 of the top 10 most influential pharmaceutical companies in Wikipedia are also among the top 10 in terms of market capitalization since 2000 and among the top 10 for market capitalization in 2017.

**Fig 3.**
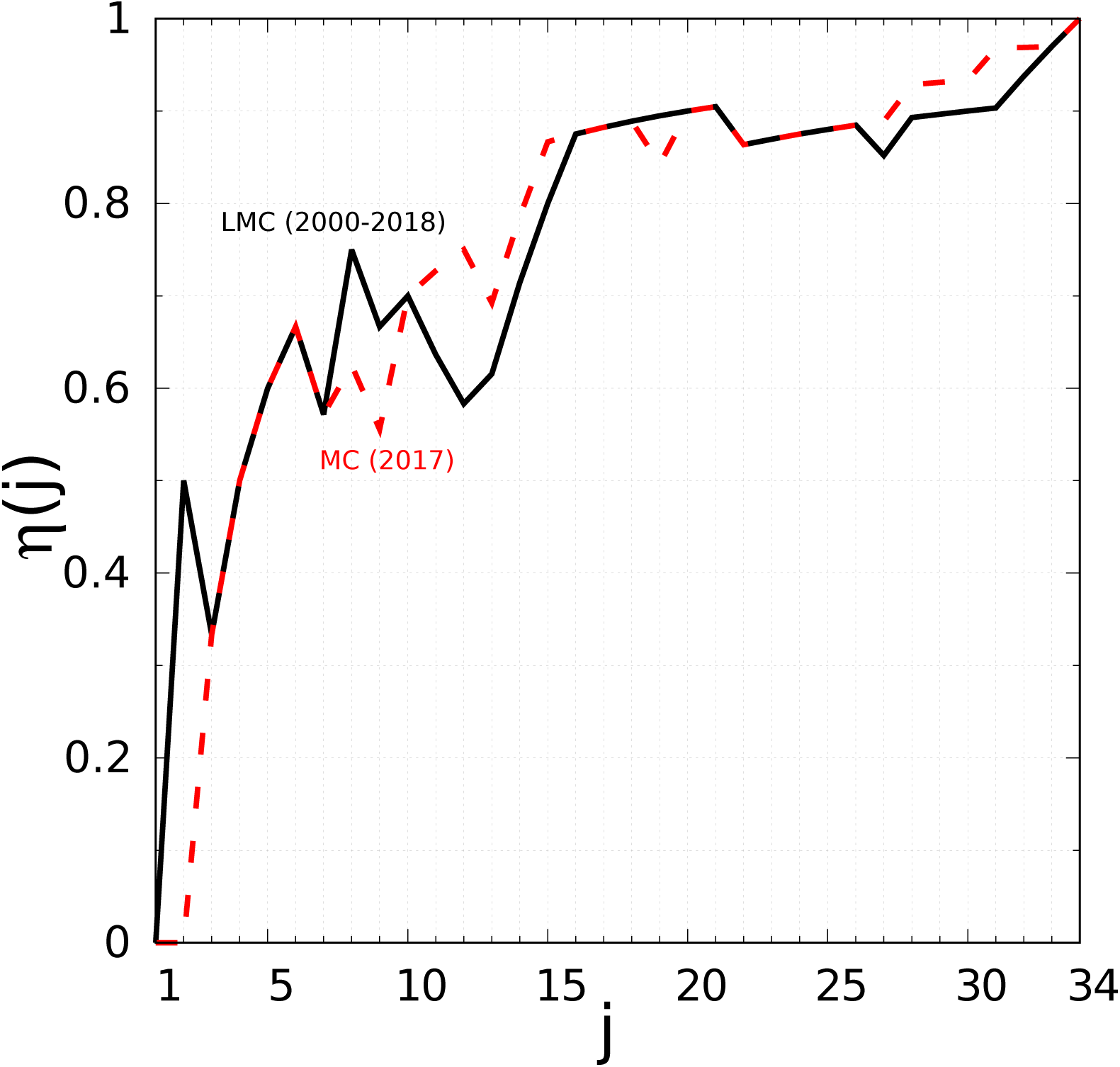
Overlap between the PageRanking of articles on pharmaceutical companies in Wikipedia and the ranking of pharmaceutical companies by market capitalization. The overlap function is *η*(*j*) = *j_ph_/j*, where *j*_*ph*_ is the number of pharmaceutical companies present in the top *j* of both lists: the PageRanking of pharmaceutical companies articles in Wikipedia (see first column of Table 1) and the list of pharmaceutical companies ranked by market capitalization in 2017 (red curve, see third column in Table 1) and ranked in terms of the largest market capitalization since 2000 (black curve, see second column in Table 1).

We have to mention that, on average, older Wikipedia articles, i.e., articles with earlier creation dates, may have accumulated more ingoing and outgoing links than newer ones and that, consequently, older Wikipedia articles should, on average, have better PageRank and CheiRank indexes. This effect is not observed among articles devoted to countries, as almost all of these articles were created during the same period, i.e., in 2001 at the very beginning of Wikipedia. However, on average, this effect is present, albeit with considerable fluctuations, among articles devoted to cancer types and pharmaceutical companies.

The distribution of the 47 articles on rare renal diseases on the PageRank-CheiRank plane is shown in Fig 4. The top three PageRank positions are taken by Alport syndrome, Bardet-Biedl syndrome and Fabry disease (*K*_*r*_ = 1, 2, 3), meaning that these are most influential rare renal diseases. The top three CheiRank positions are held by Bardet-Biedl syndrome, Kallmann syndrome and Oculocerebrorenal syndrome 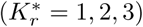, making them the most diffusive articles among the 47 articles devoted to rare renal diseases. Subsets of articles in Wikipedia such as, e.g., articles devoted to Universities [12], to infectious diseases [19] or to cancer types [20], are usually distributed around the 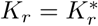 diagonal. We note that the case of rare renal diseases is different since, with the exception of the top 6 diseases in *K*_*r*_ and a few others, the rare renal diseases are located far from the 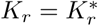 diagonal. This is corroborated by the Spearman rank correlation coefficient of *ρ* ≃ 0.54. The diseases above (below) the diagonal are more influential (diffusive) than diffusive (influential) in the Wikipedia network. We have to remember that these *K*_*r*_ and 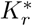 rankings are relative to the considered subset of renal diseases with high *K* and 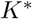 indexes, and that none of the articles devoted to these diseases are either influential or diffusive in the global Wikipedia network. Hence, for rare renal diseases, high *K*_*r*_ – low 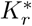 articles have more outgoing links than the average in this subset pointing toward mildly diffusive articles, e.g., articles containing lists of rare diseases. High 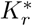– low *K*_*r*_ articles have more ingoing links than the average in this subset pointing from mildly influential articles, e.g., the top *K*_*r*_ articles on rare renal diseases.

**Fig 4.**
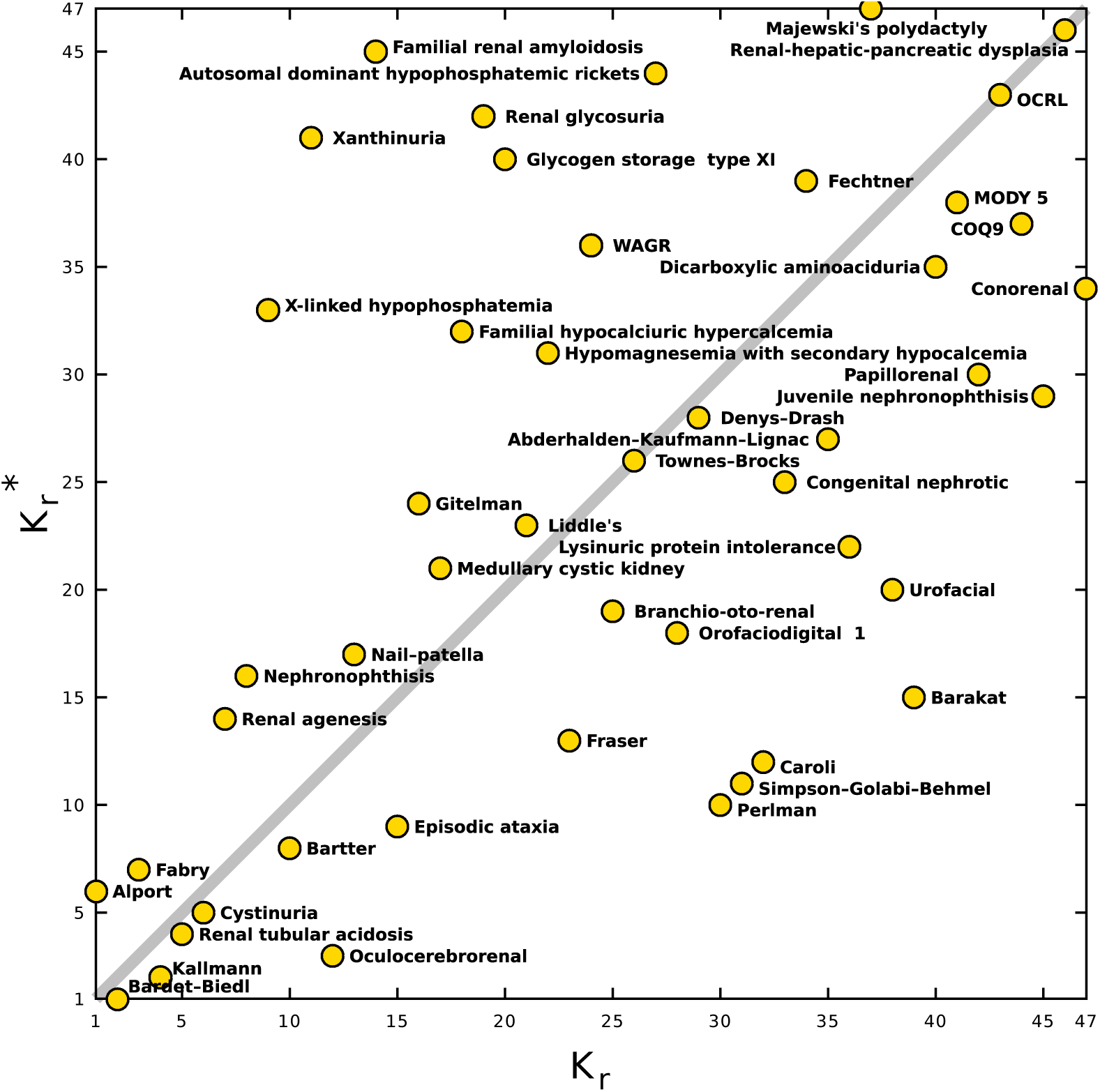
Distribution of *N*_*rd*_ = 47 articles on rare renal diseases on the plane of relative PageRank-CheiRank indexes. (*K*_*r*_, 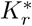);. positions in the plane are indicated by golden circles with short names for the diseases.

### Example of a reduced Google matrix *G*_R_

We will consider Wikipedia articles (nodes of interest) among the 313 selected articles (nodes) devoted to countries (*N*_*c*_ = 195), pharmaceutical companies (*N*_*ph*_ = 34), cancers (*N*_*cr*_ = 37) and rare renal diseases (*N*_*rd*_ = 47). Since the interactions between countries and cancers have been analyzed in [20], we show other interactions below, such as those between pharmaceutical companies and countries, cancers, rare renal diseases, and between rare renal diseases and countries. In Fig 5, we show the 81 *×* 81 reduced Google matrix *G*_R_ (top left panel) and its *G*_*rr*_ (top right panel), *G*_pr_ (bottom left panel), *G*_qrnd_ (bottom right panel) components associated with pharmaceutical companies and rare renal diseases. Focusing on *G*_R_ (Fig 5 top left panel), the first 34 × 34 block diagonal sub-matrix (delimited by the purple square) gives the effective directed links between pharmaceutical companies. The remaining 47 × 47 block diagonal sub-matrix (subdivided into five block diagonal sub-matrices delimited by colored squares; the color codes for the rare renal diseases are indicated in Table 2) shows the effective directed links between the rare renal diseases. The 47 × 34 (34 × 47) rectangular sub-matrix indicates the effective directed links from (to) pharmaceutical companies to (from) rare renal diseases. We clearly see from the *G*_pr_ component (Fig 5 bottom left panel) that the columns are very similar, since mathematically each column is approximately equal to the PageRank vector *P*_*r*_. Consequently, the *G*_pr_ components in their turn provides us with already known information about the global PageRank. The almost uniform rainbow pattern is due to the fact that the columns are similar and the entries are ordered by decreasing PageRank probabilities. We retrieve the fact that Wikipedia articles about top pharmaceutical companies have a higher PageRank than those concerning rare renal diseases. The 22 most influential companies have a global PageRank index from *K ∼* 10^4^ to *K ∼* 10^5^ (see Fig 1), whereas the most influential rare renal disease have a PageRank index above *K ∼* 10^5^. This is also clearly seen in the *G*_pr_ matrix picture (Fig 5, bottom left panel). The direct links are visible in the *G*_*rr*_ matrix (Fig 5 top right panel), which provides us with a picture of the adjacency matrix A. We observe that direct links are denser in the block diagonal for pharmaceutical companies than in that for rare renal diseases. In the pharmaceutical companies sector there are many direct links, with almost every Wikipedia article devoted to a pharmaceutical company pointing toward at least one other (with the exception of Alexion, BioMarin and Novo Nordisk). This is due to the very competitive nature of the industry, in which it is routine for companies to acquire drug departments from one another. Still looking at the top right panel of Fig 5, we observe that in the rare renal diseases sector, the few direct links concern rare renal diseases belonging to the same category (pixels inside the red, blue, gold cyan and green squares). The exceptions are nephronophthisis, which belongs to the ciliopathies category, and medullary cystic kidney disease, which belongs to the renal tubular diseases and metabolic diseases category; each one these diseases points directly to the other. In fact, these two diseases share similar morphological and clinical features (see, e.g., [32]), thus explaining the reciprocal direct links between them. We also observe that none of the Wikipedia articles devoted to the pharmaceutical companies with the largest market capitalization directly cite one of the rare renal diseases. This observation highlights the orphan status of these diseases. The only direct link existing between pharmaceutical companies and rare renal diseases is observed in connection with Fabry disease, which points toward Shire the manufacturer of Replagal, a dedicated drug for Fabry disease [33]. Finally, the *G*_qr_ matrix component (Fig 5 bottom right panel) indicates indirect links, some of which are purely hidden links since they do not appear in the *G*_*rr*_ matrix component (Fig 5 top right panel). The *G*_qr_ matrix component reveals many hidden links between pharmaceutical companies which are certainly due to the complex economic interconnections between these companies. The two strongest hidden links appear from Alexion to Shire and from Abbott to Johnson & Johnson. Many indirect pathways through the overall Wikipedia network contribute to these hidden links. E.g., one of the shortest paths going from Alexion to Shire is the Alexion → Ludwig N. Hantson → Shire link [34]. Indeed, Ludwig N. Hantson was appointed as CEO of Alexion Pharmaceuticals in March 2017 [35], and he was CEO of Baxalta prior its acquisition by Shire in June 2016 [36]. Furthermore, the indirect link Abbott → Advanced Medical Optics → Johnson & Johnson contributes to the Abbott → Johnson & Johnson hidden link. Indeed, Advanced Medical Optics, also known as Abbott Medical Optics, was acquired by Johnson & Johnson in February 2017 [37]. Concerning the rare renal diseases sector, the strongest hidden links among diseases belonging to the same category are: Familial hypocalciuric hypercalcemia → Gitelman syndrome, as both are hypocalciuric diseases [38]; the inverted link Gitelman syndrome → Familial hypocalciuric hypercalcemia, Juvenile nephronophthisis → Bardet–Biedl syndrome, as nephronophthisis and Bardet–Biedl syndrome are both typically cited examples of ciliopathies; Townes–Brocks syndrome → Branchio-oto-renal syndrome, as both of these syndromes imply sensorineural hearing loss [39]. Between diseases belonging to different categories, we observe the notable hidden links: Alport syndrome → Oculocerebrorenal syndrome (the inverted hidden link is also present but weaker), as both are examples of genetic causes of cataracts in childhood or early life [40]; Denys–Drash syndrome → Perlman syndrome, since both lead to a high risk of Wilms’ tumor [41, 42]. The most interesting hidden links are those that occur between pharmaceutical companies and rare renal diseases. The strongest of these is Alexion → Fabry disease. Indeed, in May 2015, Alexion Pharmaceuticals acquired Synageva BioPharma Corp., a company dedicated to rare diseases and to Fabry disease in particular [43]. We also observe hidden links, although somewhat weaker, from BioMarin and Vertex to Fabry disease. Here, it is not easy to find indirect pathways through Wikipedia from Alexion, BioMarin, or Vertex to Fabry disease unlike the case for hidden links exclusively between the cited pharmaceutical companies or exclusively between rare renal diseases.

**Fig 5.**
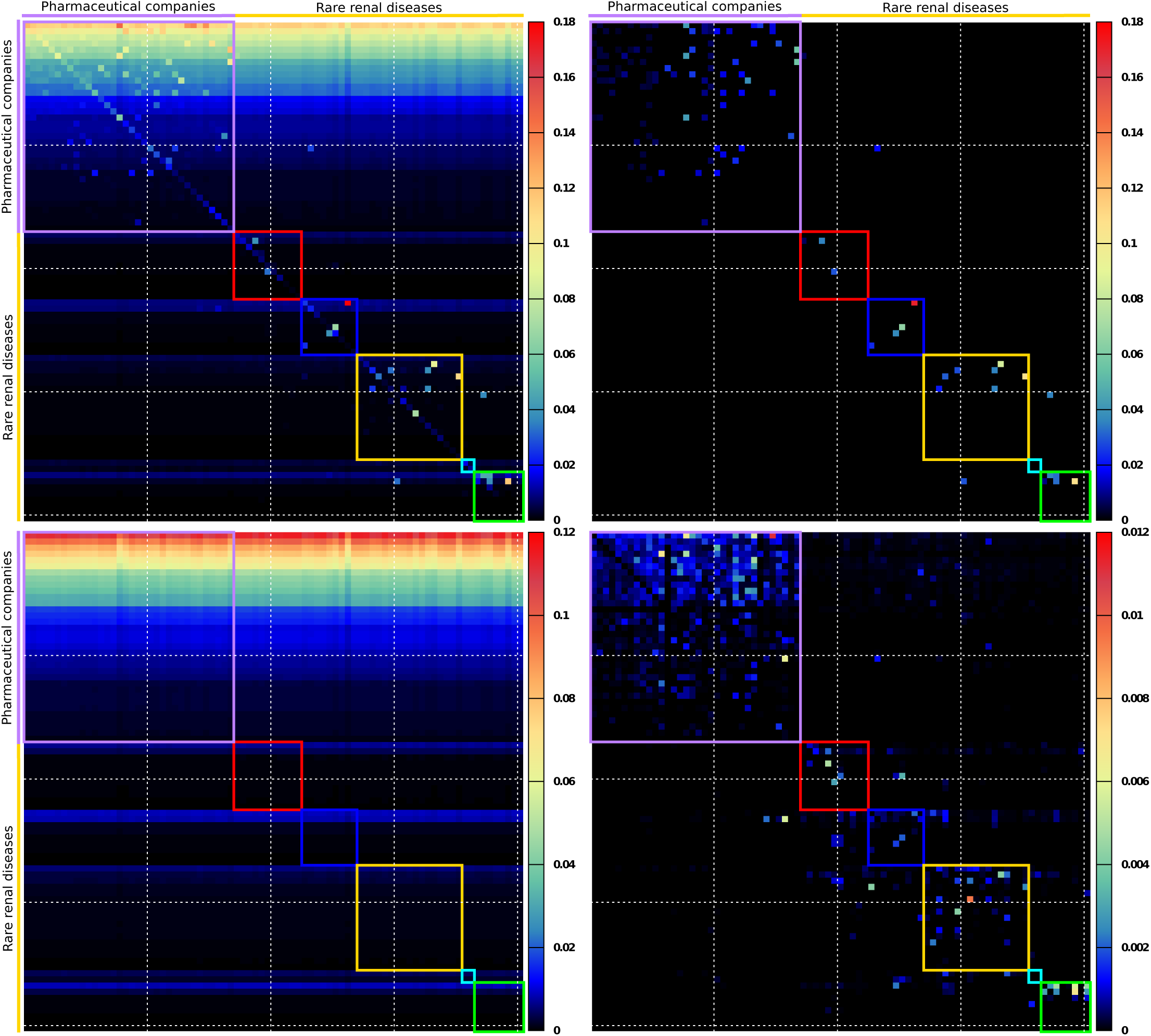
Reduced Google matrix *G*_R_ of pharmaceutical companies and rare renal diseases. We show the reduced Google matrix *G*_R_ (top left panel) and its three components *G*_*rr*_ (top right panel), *G*_pr_ (bottom left panel), and *G*_qrnd_ (bottom right panel). Each “pixel” represents a matrix entry with the amplitude given by a color. Color bars give the corresponds between matrix entry amplitudes and colors. For each 81 × 81 matrix the first 34 entries correspond to pharmaceutical companies (ordered as in Table 1) and the other 47 entries correspond to rare renal diseases (ordered by categories then by PageRank order inside each category, see Table 2). The first 34 × 34 block diagonal sub-matrix (purple square) corresponds to directed interactions between pharmaceutical companies. The other five smallest block diagonal sub-matrices correspond to directed interactions between rare renal diseases belonging to one of the five categories defined in Table 2. The colors of the squares correspond to color categories given in Table 2. For the sake of visibility horizontal and vertical white dashed lines are drawn after every 20 entries.

### Reduced network structures

As the *G*_pr_ matrix component of the *N_r_ × N*_*r*_ reduced Google matrix *G*_R_ gives trivial results that are already known from a direct PageRank analysis of the *N × N* global Google matrix, we use the *G*_*rr*_ + *G*_qr_ matrix component to infer an effective reduced network between the *N*_*r*_ nodes of interest.

First, we build the reduced network of interactions between the *N*_*r*_ = *N*_*ph*_ + *N*_*c*_ nodes of interest constituted by the *N*_*ph*_ = 34 pharmaceutical companies and the *N*_*c*_ = 195 countries (see Fig 6). The rules for the construction of the reduced network are described in the caption to Fig 6. At the first level, we take the top 5 companies in the PageRank list (Table 1) and trace the links with the largest column matrix elements in *G*_*rr*_ + *G*_qr_ from a company to two countries and two other companies. The process is then repeated to other levels from the newly added companies. The process is stopped when no new companies can be added. We obtain a compact reduced network of 15 pharmaceutical companies from the 34 initially selected, and 12 of these constitute the top 12 of the PageRank list (Table 1). The 3 remaining companies are nevertheless in the top 9 of pharmaceutical companies with the largest market capitalization since 2000.

**Fig 6.**
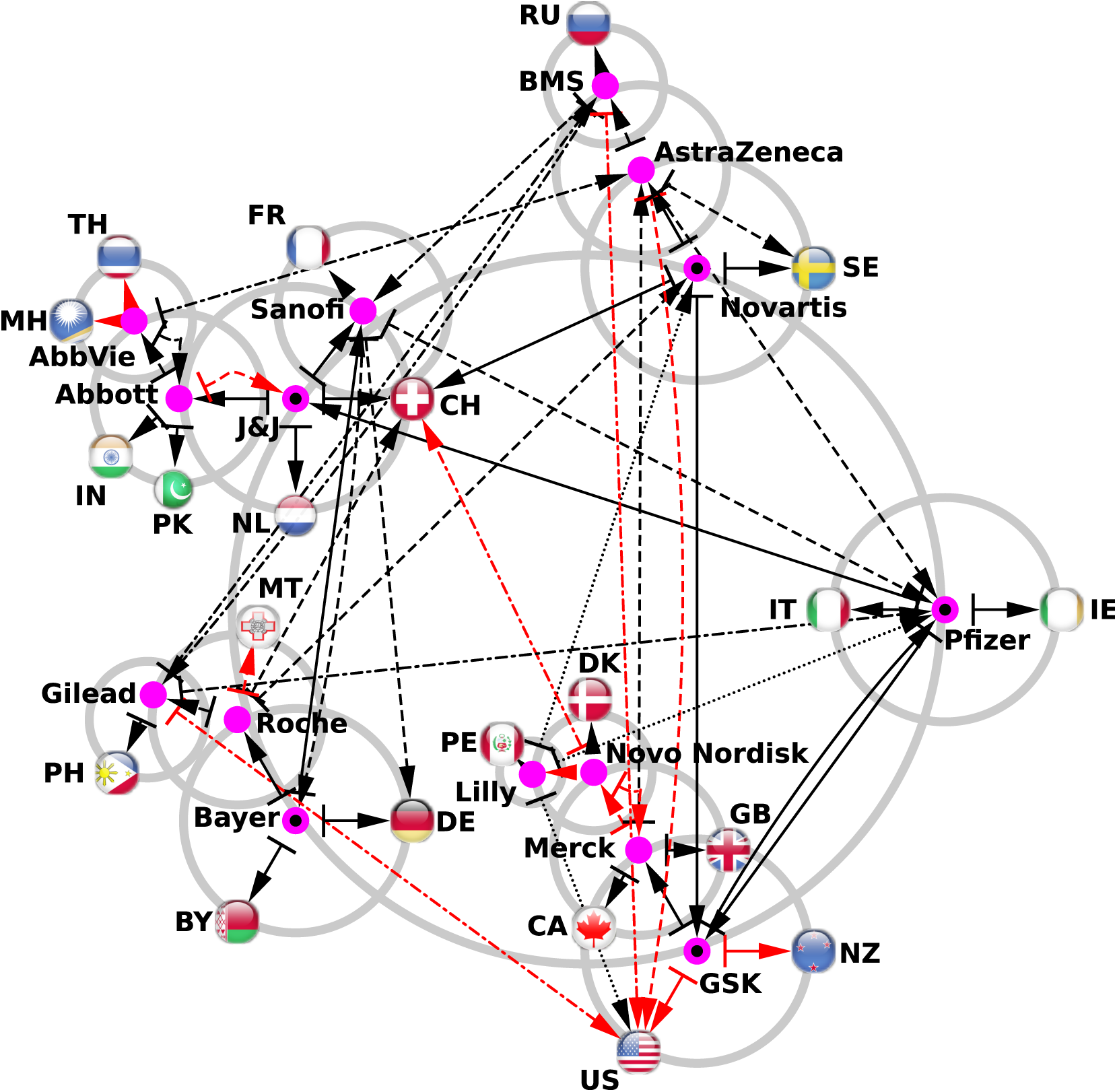
Reduced network of pharmaceutical companies with the addition of their best connected countries. We consider the first five pharmaceutical companies from the Wikipedia PageRank list: Pfizer, GSK, Bayer, J&J, and Novartis (see Table 1). Each one of these companies are represented by purple circles 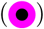 placed along the main grey circle (the grey circle with the largest radius). From these five most influential pharmaceuticals in Wikipedia, we determine the two best connected companies, i.e., for a pharmaceutical company *ph*, we determine the two companies *ph*_1_ and *ph*_2_ giving the highest 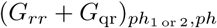 values. If not already present in the network, we add these best connected companies along secondary circles centered on the previous companies. The newly added companies are represented by purple circles 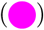. Also from the initial five pharmaceutical companies we determine the two best connected countries, i.e., for a company *ph*, we determine the two countries *c*_1_ and *c*_2_ giving the highest 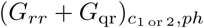 values. From the newly added pharmaceutical of the first iteration, we determine the two best connected pharmaceutical companies and the two best connected countries. This constitutes the second iteration, an so on. At the fourth iteration of this process, no new pharmaceutical companies are added, and consequently the network construction process stops. The links obtained at the first iteration are represented by solid line arrows, at the second iteration by dashed line arrows, at the third iteration by dashed-dotted line arrows, and at the fourth iteration by dotted line arrows. Black color arrows correspond to links existing in the adjacency matrix (direct hyperlinks in Wikipedia), and red color arrows are purely hidden links absent from the adjacency matrix but present in *G*_qr_ component of the reduced Google matrix *G*_R_. The obtained network is drawn with the Cytoscape software [44]. Countries are marked by their ISO 3166-1 alpha-2 codes.

By following direct links (black arrows) in this network, i.e., hyperlinks from Wikipedia articles, we see, e.g., that for Bayer, the closest companies are Sanofi and Roche and the two country friends are Germany, where its main office is located, and Belarus. These two links are direct links from the Bayer page in Wikipedia. Here, Belarus is cited as an example of a country from the Commonwealth of Independent States (CIS) in which the over-the-counter drug business was developing in early 2000s. To respond to this need, Bayer acquired Sagmel, Inc., a company already present in the CIS, in June 2008 [45, 46]. For Pfizer, the two company friends are GlaxoSmithKline and Johnson & Johnson, while the two country friends are Italy and Ireland. All these direct links are easy to explain by examining the hyperlinks in Wikipedia articles. Direct links between companies indicates major transactions between them that are reported in the dedicated Wikipedia articles. These direct links give a compact picture of the relationships between pharmaceutical companies. The non-obvious hidden links (red arrows in Fig 6) are potentially more interesting. Here, most of the hidden links point to the US, highlighting the fact that most of the pharmaceutical companies are American.

The reduced network of interactions between the *N*_*ph*_ = 34 pharmaceutical companies and the *N*_*cr*_ = 37 cancer types is shown in Fig 7. At each level, it shows the two closest company friends and two cancer types. The construction rules are the same as in Fig 6. The structure of the reduced network between pharmaceutical companies remains the same as in Fig 6. The reduced network indicates the strongest links from a company to the related types of cancer. We observe a clear polarization toward breast cancer, since 10 of the 15 pharmaceutical companies preferentially point to breast cancer, the Wikipedia article for which is the second most influential article among the 37 articles devoted to cancer types [20]. The second and third most connected cancer types are leukemia and lung cancer, which are preferentially pointed to by 4 and 3 pharmaceutical companies, respectively. Although the list (lung cancer, breast cancer, and leukemia) contains the 3 most influential cancer types in Wikipedia [20], we observe the particular interest in breast cancer on the part of pharmaceutical companies. More specifically, Pfizer and GlaxoSmithKline are most strongly linked to breast cancer and leukemia, while Bayer is most strongly linked to breast and thyroid cancers. In the case of Johnson & Johnson, the strongest links are to leukemia and ovarian cancers. For Novartis, the strongest links are with melanoma and breast cancers. Links to cancers for other companies are also clearly visible in Fig 7. We argue that the REGOMAX approach allows us to determine, from Wikipedia, the main orientations of pharmaceutical companies in their treatment of cancers. From Fig 7, we also see that in addition to direct links (black arrows), the indirect links (red arrows) also play a very important role.

**Fig 7.**
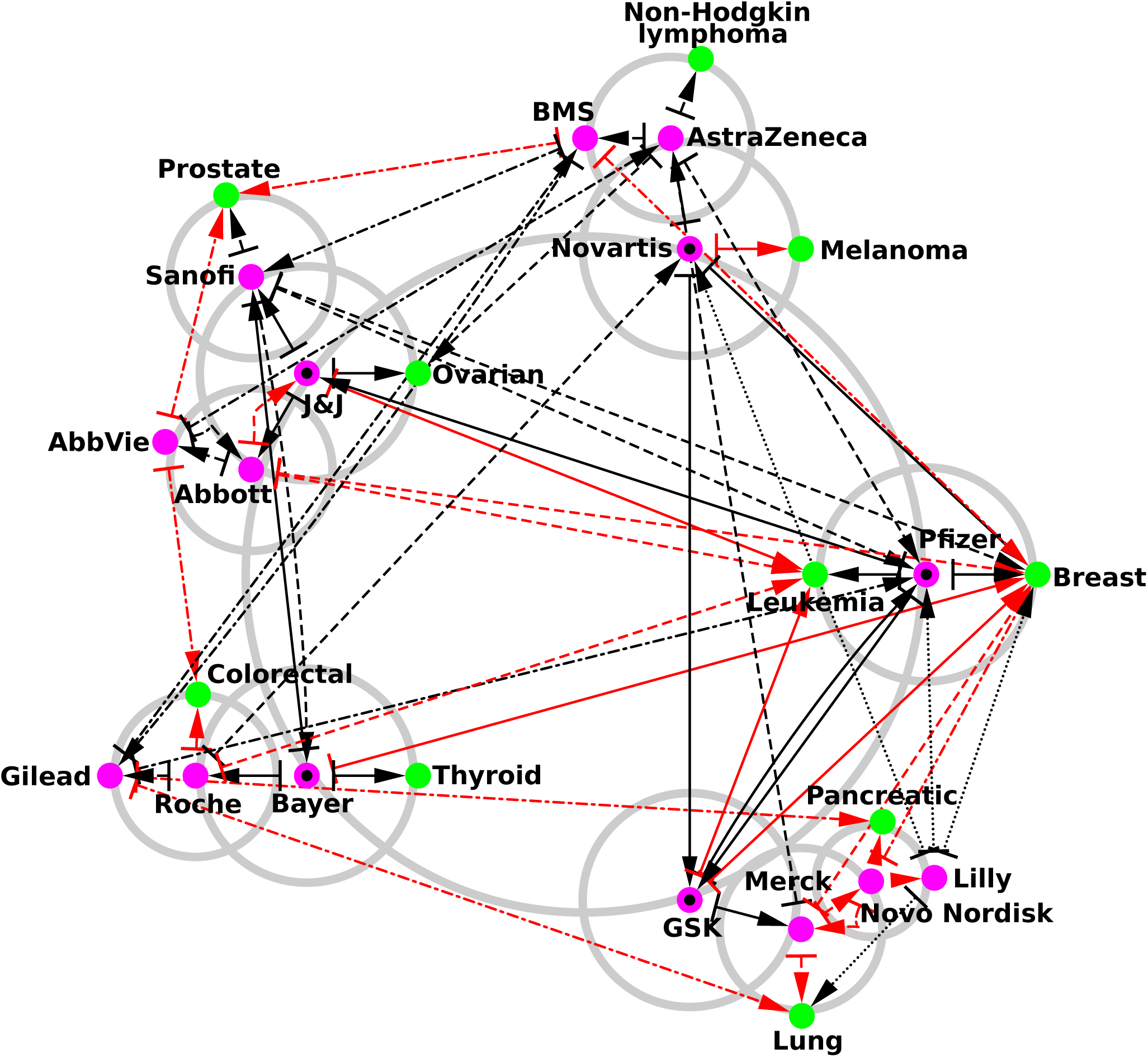
Reduced network of pharmaceutical companies with the addition of their best connected cancers. The construction algorithm is the same as the one used to generate Fig 6 excepting that we replace at each iteration the two best connected countries by the two best connected cancers. Pharmaceutical companies are represented by purple circles 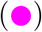 and cancers by green circles 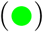.

The reduced network of pharmaceutical companies and rare renal diseases is shown in Fig 8. Its construction rules are the same as for Figs 6 & 7. Again, the network structure of the pharmaceutical companies remains the same as in Figs 6 & 7. Since 15 pharmaceutical companies are present, 30 preferentially connected rare renal diseases could have potentially emerged from this reduced network, but in fact only 10 rare renal diseases are present. From this network, we directly see the main orientations of each company toward specific rare renal diseases. We observe a strong polarization toward Fabry disease (*K*_*r*_ = 3 in Table 2) since 12 of the 15 pharmaceutical companies point to this disease. The second and third most connected rare renal diseases are Alport syndrome (*K*_*r*_ = 1 in Table 2) with 5 ingoing links and Kallmann syndrome (*K*_*r*_ = 4 in Table 2) with 3 ingoing links. Looking in more detail, Pfizer has the strongest links to Kallmann syndrome and Fabry disease. GlaxoSmithKline is linked to Fabry disease and medullary cystic kidney disease, while Bayer is oriented toward renal agenesis and familial renal amyloidosis disease. For Johnson & Johnson, the main orientations are toward Alport syndrome and Fabry disease. Novartis is mainly linked to renal tubular acidosis and Fabry disease. We observe that none of the articles devoted to the pharmaceutical companies with the largest market capitalization cite any of the rare renal diseases. All the connections from pharmaceutical companies to rare renal diseases are therefore indirect hidden links (red arrows in Fig 8), thereby indeed confirming the effective status of orphan diseases.

**Fig 8.**
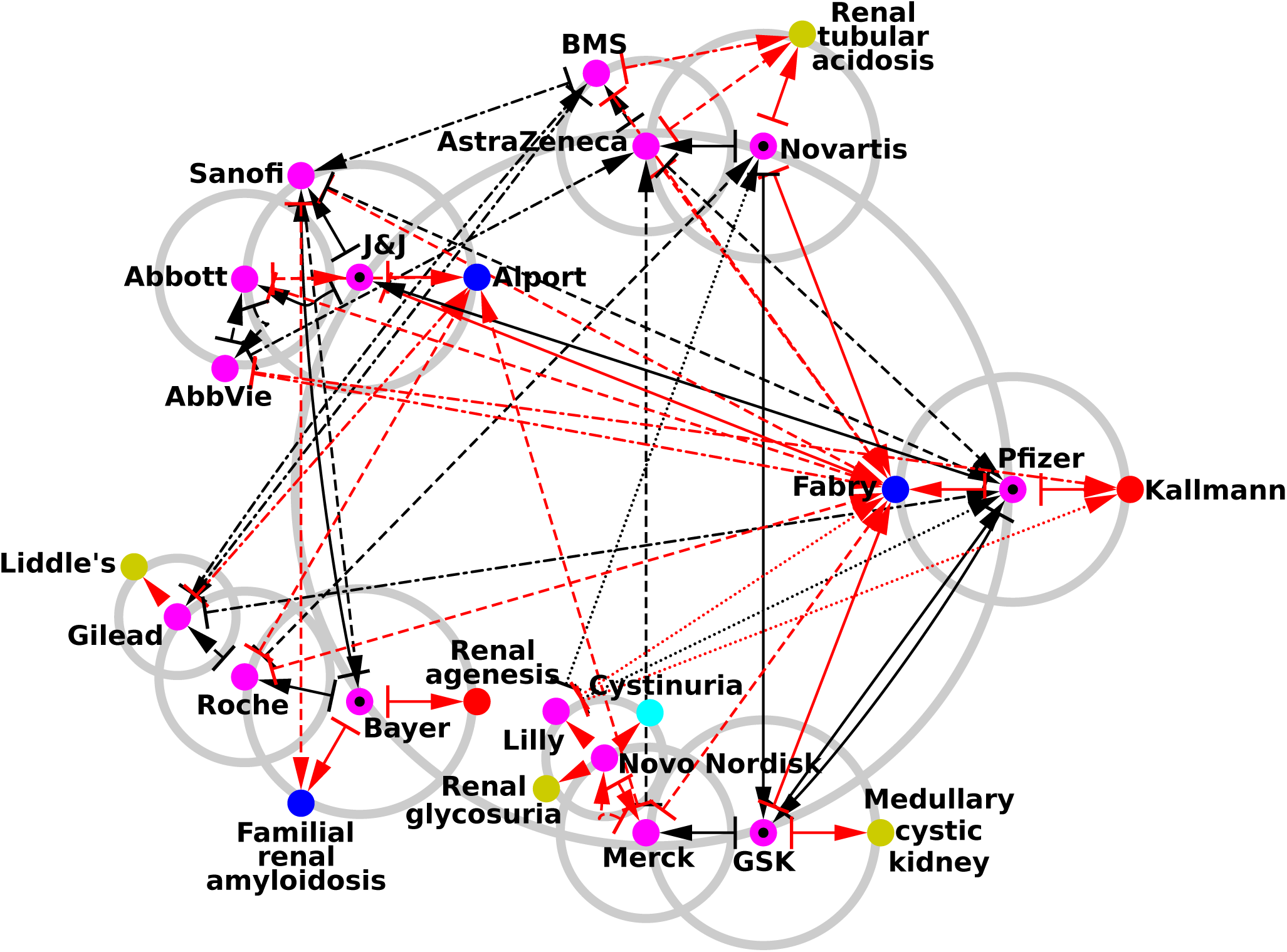
Reduced network of pharmaceutical companies with the addition of their best connected rare renal diseases. The construction algorithm is the same as the one used to generate Fig 6 excepting that we replace at each iteration the two best connected countries by the two best connected rare renal diseases. Pharmaceutical companies are represented by purple circles 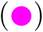 and rare renal diseases by red circles 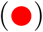 for congenital abnormalities of the kidney and urinary tract, blue circles 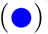 for glomerular diseases, gold circles 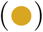 for renal tubular diseases and metabolic diseases, cyan circles 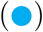 for nephrolithiasis, and green circles 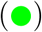 for ciliopathies.

We also present the friendship network between companies and infectious diseases studied in [19] in SupInfo Fig S2.

### Sensitivity of countries to pharmaceutical companies within Wikipedia

To see in Wikipedia, the influence of a specific pharmaceutical company on different countries we use the PageRank sensitivity *D* described in the Section Reduced Google matrix.

The PageRank sensitivity of an article devoted to country *c* to a specific pharmaceutical company article *ph* is determined by *D*(*ph→c, c*). This quantity measures the relative change of the country *c* article’s PageRank probability caused by the increase in the intensity of the transition probability from the pharmaceutical company *ph* article to the country *c* article. In other words, the PageRank sensitivity *D*(*ph→c, c*) measures what happens to the influence of the country *c* article in Wikipedia when there is increased interest in the country *c* article resulting from the pharmaceutical company *ph* article. In the following, the expression *sensitivity of item A to item B* refers to the *PageRank sensitivity of the item A article to the item B article*. In Fig 9, we show the sensitivity of world countries to two companies, Pfizer and Bayer. The two most sensitive countries for Pfizer are Ireland and Italy. This observation correlates with the strongest direct links shown in Fig 6. Italy is present here for historical reasons as the calcium citrate used to produce citric acid was supplied to Pfizer by Italy until a shortage caused by World War I occurred, thus forcing Pfizer’s chemists to develop fermentation technology to obtain citric acid from sugar by using a fungus. This technology was then used on a large scale to produce antibiotic penicillin during World War II [48]. In 2016, Pfizer attempted to acquire Allergan, an Irish–tax registered pharmaceutical company, in order to create the world’s largest drugmaker and relocate its headquarters in Ireland to reduce taxes. The deal was called off after the US Treasury adopted new anti-tax inversion rules [49]. The next countries most impacted by Pfizer impacted countries are Canada, Australia, and New Zealand. There is no direct link between Pfizer and these countries. Among the shortest indirect paths, we have, e.g., Pfizer *→* Terre Haute *→* Canada; Pfizer *→* Wyeth *→* Australia; and Pfizer → Helen Clark → Judicial Committee of the Privy Council → New Zealand. Pfizer has been present in Canada since the fifties, and in 2012, Pfizer’s Canadian division was recognized as one of the 15 best places to work in Canada [50]. In 2009, Pfizer acquired Wyeth which supplied a pneumococcal vaccine approved for young children in Australia [51]. Wikileaks revealed that in 1990, Pfizer was lobbying in the US against New Zealand, whose drug-buying rules it considered to be restrictive; Helen Clark was at that time New Zealand Health Minister [52]. For Bayer the most sensitive countries are Belarus, Ukraine and Germany. Indeed, the article on Bayer has direct links to Belarus and Ukraine. There is also direct link to Germany and many indirect links to it since the company is located in Germany. It is possible that Germany is less sensitive to Bayer than Ukraine and Belarus are since its PageRank probability is significantly higher. We find again here Belarus and Ukraine, which are members of the CIS, as is explained the in Section Reduced network structures. India also appears as it hosts a part of Bayer Business Services [53].

**Fig 9.**
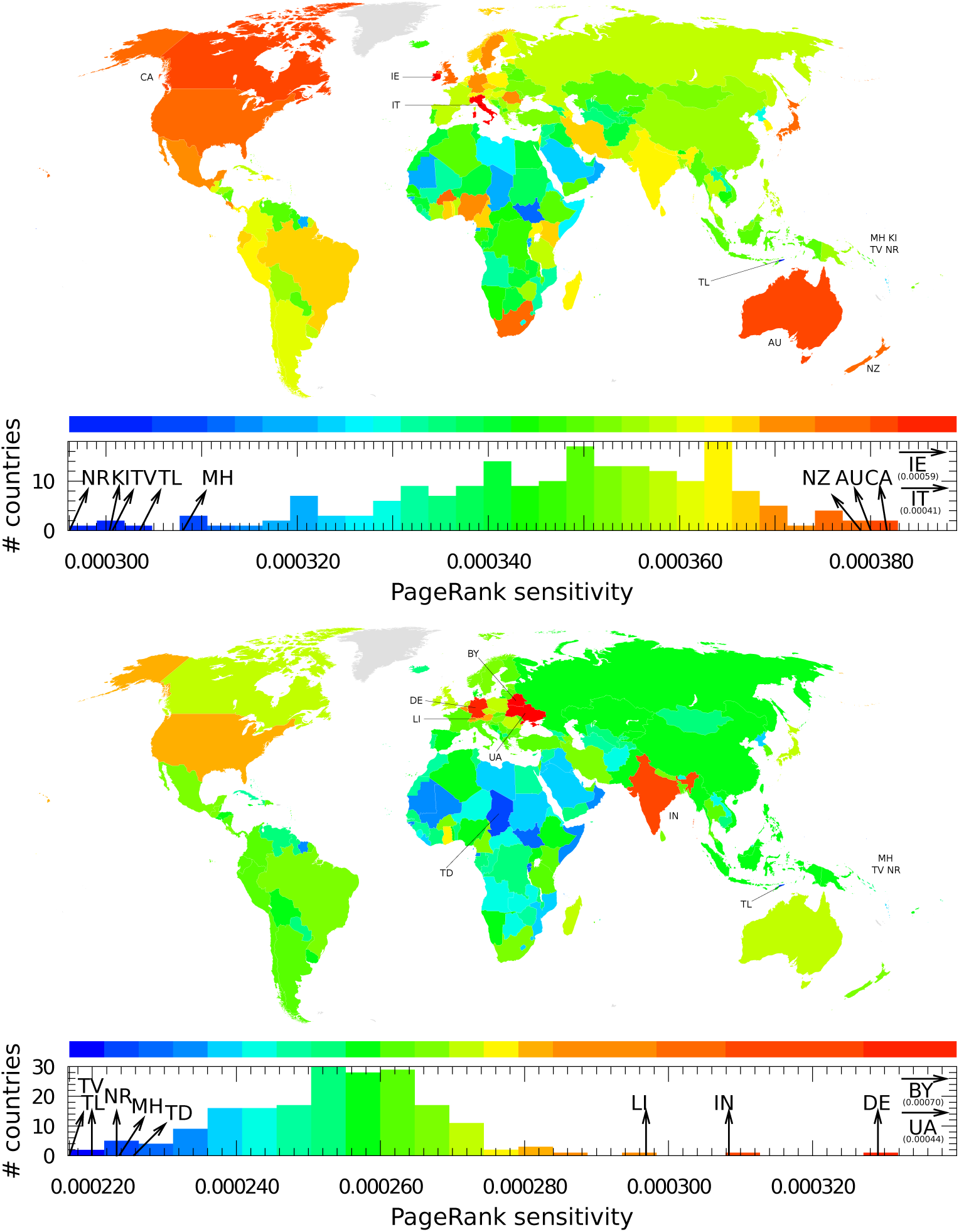
Sensitivity of countries to the Pfizer company (top panel) and the Bayer company (bottom panel). A country *c* is colored according to its diagonal PageRank sensitivity *D*(*ph→c, c*), where *ph* is the pharmaceutical company. Color categories are obtained using the Jenks natural breaks classification method [47].

### Sensitivity of countries to rare renal diseases within Wikipedia

The PageRank sensitivity of a country *c* article to a specific rare renal disease *rd* is determined by *D*(*rd→c, c*). In Fig 10, we present the PageRank sensitivity of countries to Kallmann syndrome and Bardet–Biedl syndrome. For Kallmann syndrome, the most sensitive countries are Switzerland, Germany and Spain. From Kallmann syndrome, many paths converge toward Germany. We have, e.g., the direct link Kallmann syndrome → Germany, and, e.g., the indirect link Kallmann syndrome → Franz Josef Kallmann → Germany. Franz Josef Kallmann who gave the first description of this disease, was a German-born geneticist. The Wikipedia article devoted to Kallmann syndrome points to Switzerland through a direct link and an indirect link: Kallmann syndrome → Lausanne University Hospital → Switzerland. The Lausanne University Hospital is cited as one of the main research facilities working on this disease [54]. The Spanish doctor Aureliano Maestre de San Juan also identified the link between anosmia and hypogonadism (Kallmann syndrome → Aureliano Maestre de San Juan → Spain). Country sensitivity appears to be due to the combination of direct and indirect links captured by the REGOMAX algorithm.

**Fig 10.**
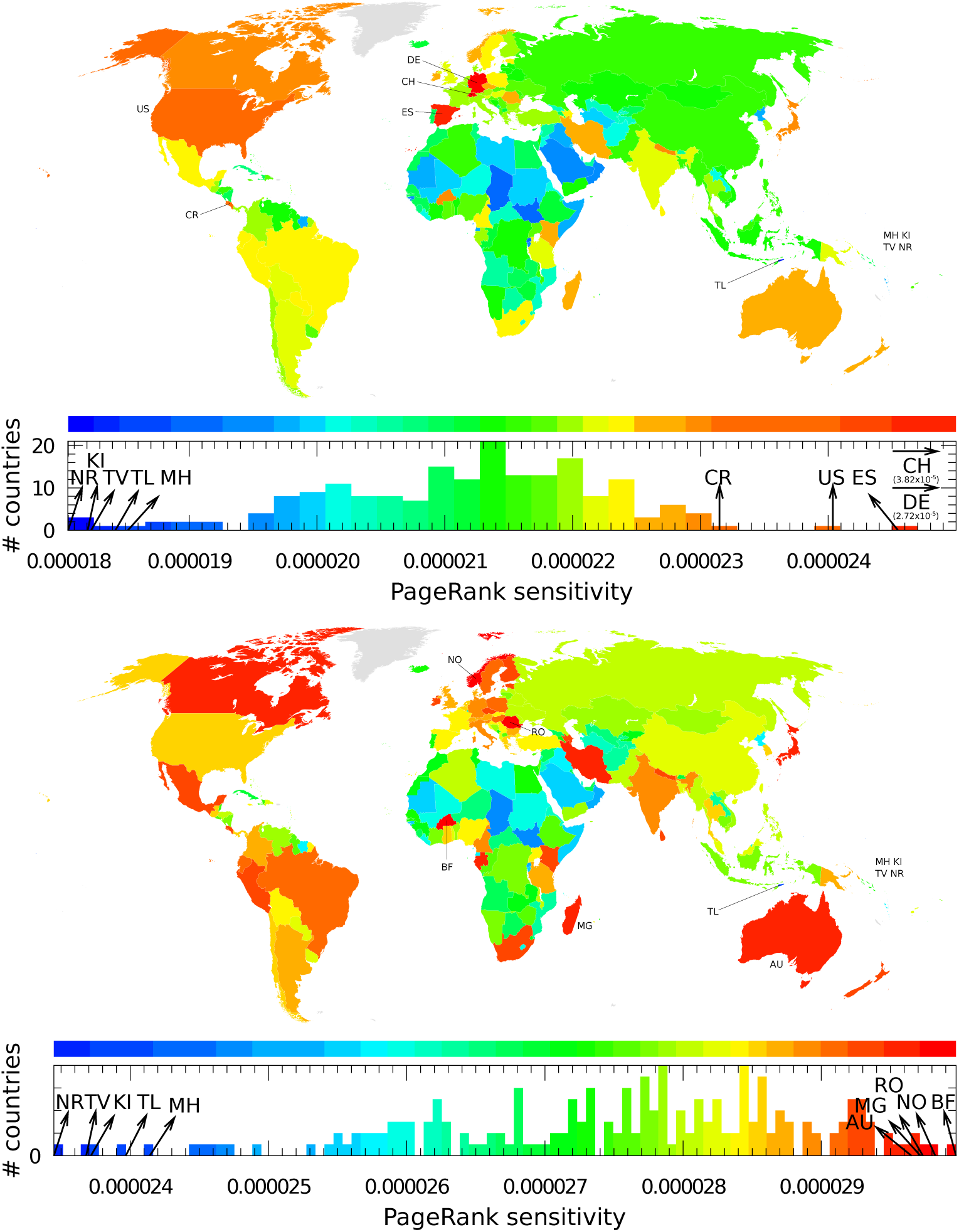
**Sensitivity of countries to rare renal diseases;** here to Kallmann syndrome (top panel) and to Bardet–Biedl syndrome (bottom panel). A country *c* is colored according to its diagonal PageRank sensitivity *D*(*rd→c, c*), where *rd* is the rare renal disease. Color categories are obtained using the Jenks natural breaks classification method [47].

In the case of Bardet–Biedl syndrome, the most sensitive countries are Burkina Faso, Norway and Romania. Here, there are no direct links to these countries from the article on Bardet–Biedl syndrome [55]. The indirect link to Romania appears because Arthur Biedl was born in what is now Romania. Indirect paths leading to the other most sensitive countries are difficult to find.

We also present the sensitivity of pharmaceutical companies to cancers and rare renal disease, respectively, in Fig S3 and Fig S4 in the Section Supplementary Information.

## Discussion

We used the reduced Google matrix (REGOMAX) algorithm for an analysis of the English Wikipedia network with more than 5 million articles. The analysis focused on 195 world countries, the 34 largest biotechnology and pharmaceutical companies, 47 rare renal diseases and 37 types of cancer. The algorithm made it possible to construct the reduced Google matrix of these entries, while also taking account of the direct and indirect links between them. While the direct links were directly present in the global Wikipedia network, the indirect links were obtained with REGOMAX by summing the contributions between entries from all the pathways connecting them via the global network. Using the reduced Google matrix, we were able to determine the interaction networks between companies and countries, companies and rare renal diseases, and companies and cancers. The sensitivity of PageRank probabilities enabled us to identify the influence in Wikipedia of specific companies on countries. This approach also revealed the sensitivity of countries to specific rare renal diseases. We identified the most influential pharmaceutical companies and showed that the top PageRank positions are occupied by companies which are far from heading the market capitalization list.

We argue that improving Wikipedia articles about specific pharmaceutical companies can increase their worldwide visibility without significant additional investments. Our study shows that the knowledge accumulated in Wikipedia can be efficiently analyzed using the REGOMAX algorithm, which determines the effective interactions between specific Wikipedia articles that are of interest for researchers.

## Acknowledgments

This work was partially supported by the Programme Investissements d’Avenir ANR-11-IDEX-0002-02, reference ANR-10-LABX-0037-NEXT (THETRACOM project). It was also partially supported the Programme Investissements d’Avenir ANR-15-IDEX-0003, ISITE-BFC (GNETWORKS project) and by the Bourgogne Franche-Comté region (APEX project).

## Supplementary Information

In Fig S1 we present in the relative PageRank *K*_*r*_ – CheiRank 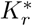 indexes plane the set of selected Wikipedia articles described in the Section Datasets. In Fig S2 we show the friendship network of interactions between pharmaceutical companies and infectious diseases studied in [19].

We also show that the REGOMAX analysis allows us to determine the inter-sensitivity of pharmaceutical companies to cancers and rare renal diseases. We show the sensitivity of 34 pharmaceutical companies to 37 cancers and 47 rare renal diseases (and vice versa) in Fig S3 and Fig S4 respectively. We note that the very strong matrix element in Fig S4 (red square) between Shire and Fabry disease appears due to the drug Replagal which is a treatment for this disease and is produced by Shire.

**Fig S1.**
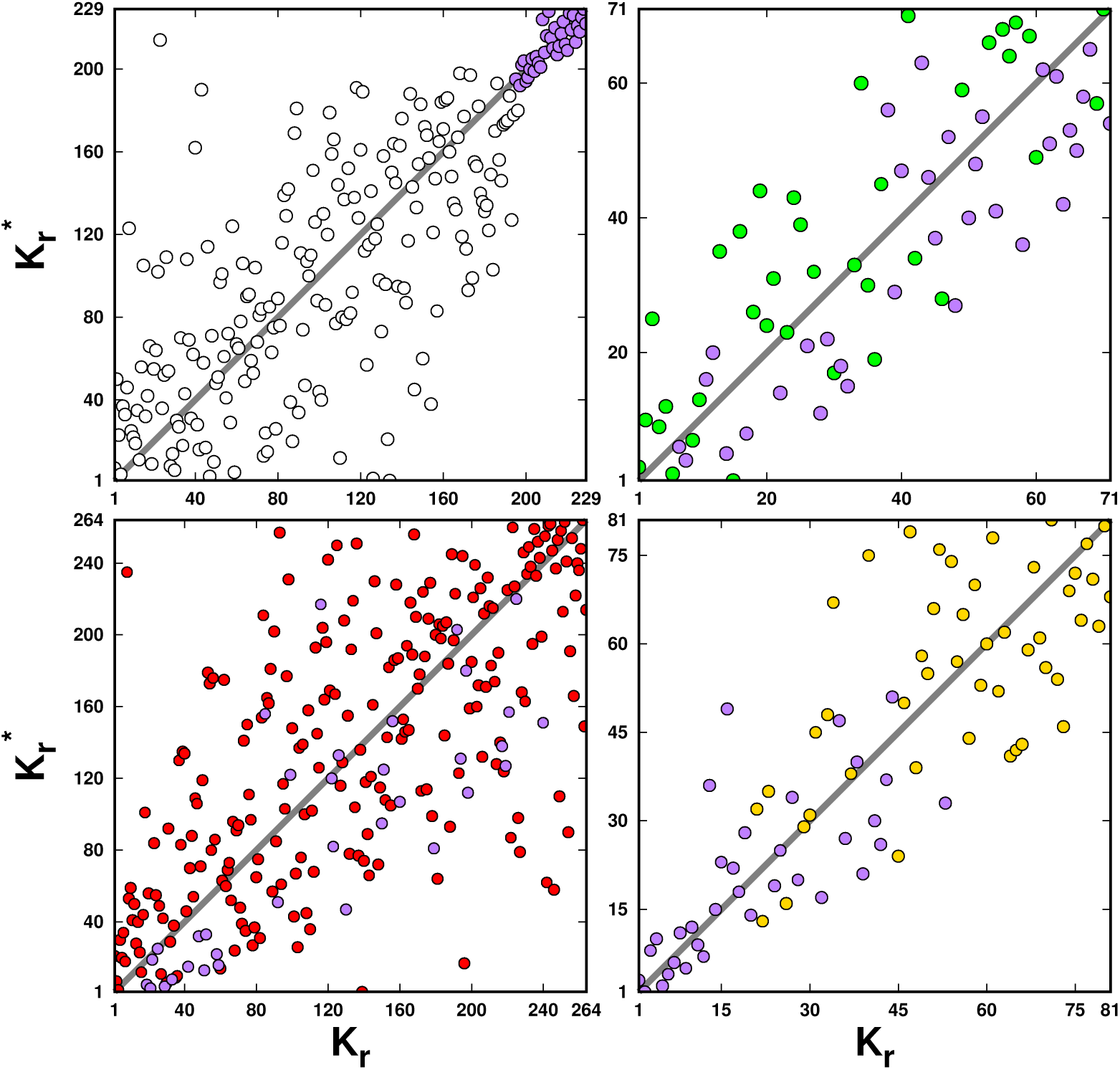
Distribution of the May 2017 English Wikipedia articles in the relative PageRank *K*_*r*_ – CheiRank 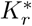 indexes plane for pharmaceutical companies and countries (top left panel), cancer types (top right panel), infectious diseases (bottom left panel), and rare renal diseases (bottom right panel). The *N*_*ph*_ = 34 pharmaceutical companies are represented by purple circles 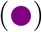, the *N*_*c*_ = 195 countries by white circles 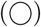, the *N*_*cr*_ = 37 cancer types by green circles 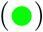, the *N*_*d*_ = 230 infectious diseases by red circles 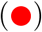, and the *N*_*rd*_ = 47 rare renal diseases by gold circles 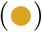.

**Fig S2.**
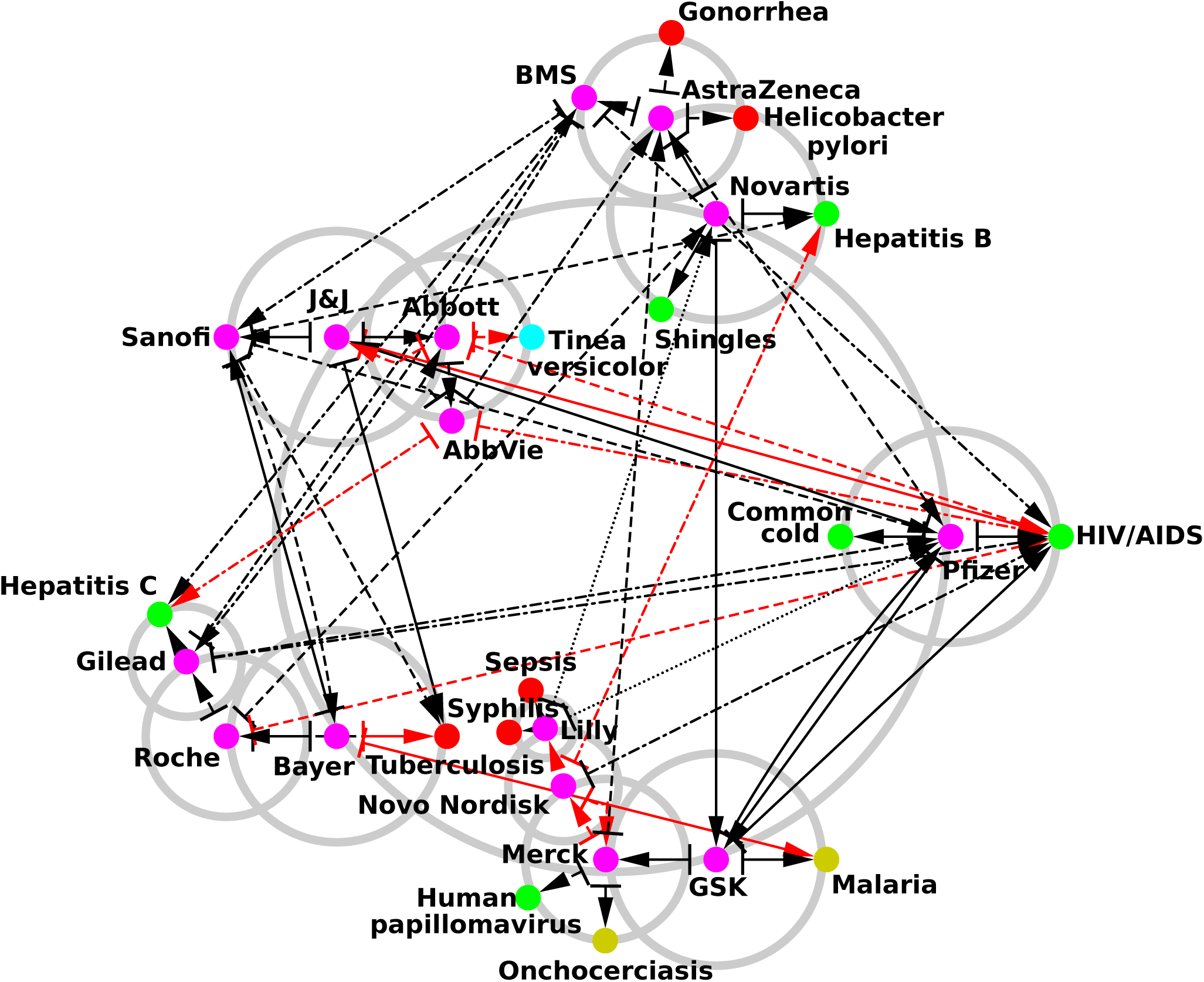
Reduced network of pharmaceutical companies with the addition of their best connected infectious diseases. The construction algorithm is the same as the one used to generate Fig 6 excepting that we replace at each iteration the two best connected countries by the two best connected infectious diseases. Pharmaceutical companies are represented by purple circles 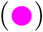 and infectious diseases by red circles 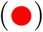 for bacterial type diseases, green circles 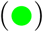 for viral type diseases, gold circles 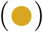 for parasitic type diseases, and cyan circles 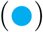 for fungal type diseases. The list of infectious diseases is available in [19].

**Fig S3.**
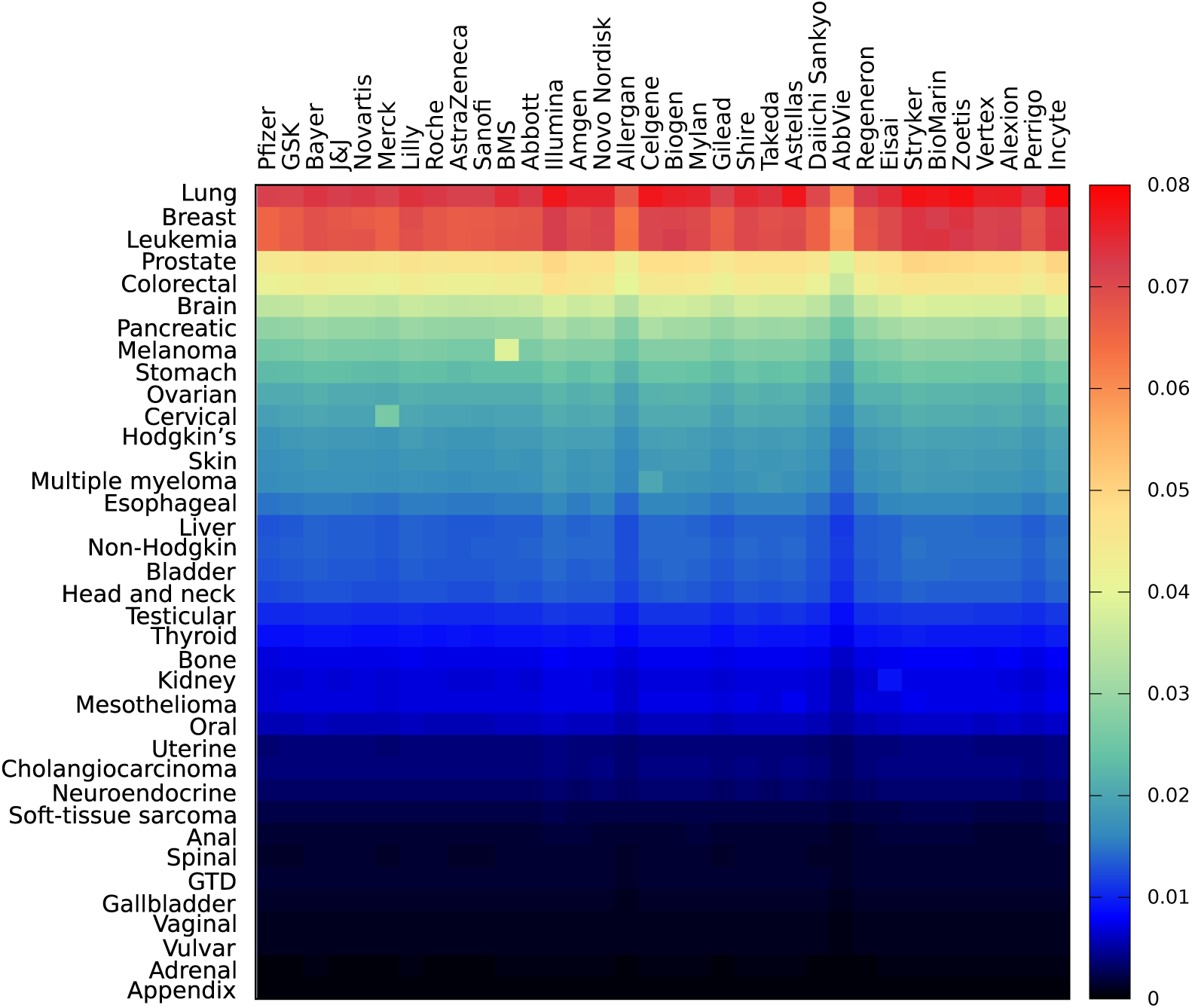
Sensitivity of pharmaceutical companies to cancers. Horizontal (vertical) entries represents pharmaceutical companies (cancer types). The acronym GTD stands for gestational trophoblastic disease.

**Fig S4.**
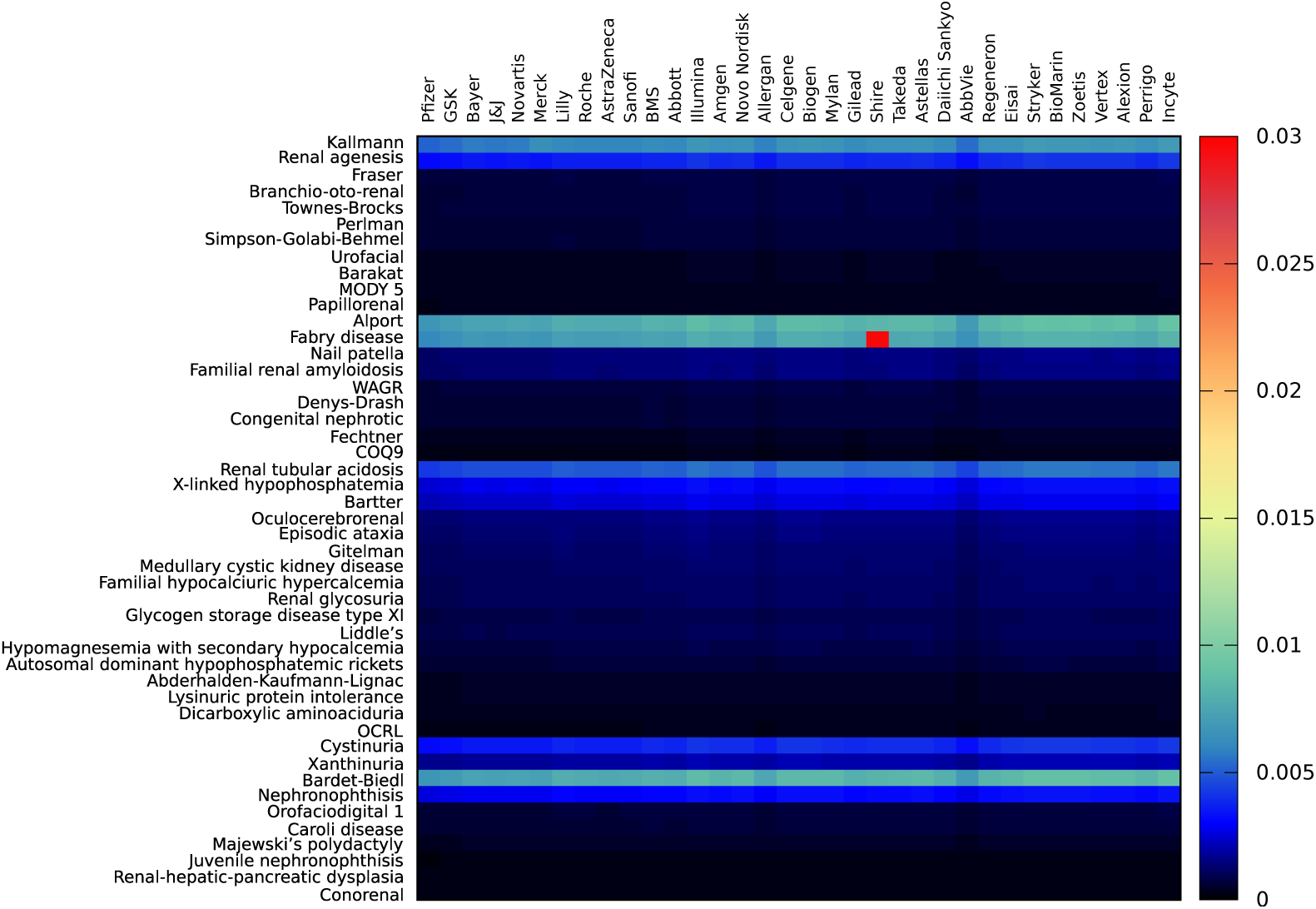
Sensitivity of pharmaceutical companies to rare renal diseases. Horizontal (vertical) entries represents pharmaceutical companies (rare renal diseases).

## Notes

http://perso.utinam.cnrs.fr/~lages/datasets/WikiPharma

